# Statin-induced Mitochondrial Coenzyme Q Deficiency Alters Mitochondrial Redox Homeostasis and Bioenergetic Function in Astrocytes

**DOI:** 10.64898/2026.06.10.731318

**Authors:** Krzysztof Wojcicki, Lukasz Galganski, Adrianna Budzinska, Grzegorz Figura, Wojciech Pijanowski, Wieslawa Jarmuszkiewicz

## Abstract

Statins, widely used cholesterol-lowering drugs, inhibit the mevalonate pathway and reduce coenzyme Q (CoQ) biosynthesis, potentially impairing mitochondrial function. Because astrocytes are essential for maintaining brain redox homeostasis, statin-induced mitochondrial dysfunction in these cells may contribute to central nervous system pathology. Therefore, this study examined the effects of a 6-day statin exposure on mitochondrial bioenergetics in rat astrocytes, focusing on mitochondrial CoQ (mtCoQ) deficiency. Treatment with 200 nM atorvastatin or simvastatin decreased the total mtCoQ pool (mtCoQ9 + mtCoQ10) by 30%–35% and the antioxidant pool (mtCoQH_2_) by 40% but did not alter the levels of mitochondrial antioxidant proteins, including superoxide dismutase 2 and uncoupling proteins. Mitochondria of statin-treated astrocytes showed decreased respiratory activity, membrane potential, and ATP synthesis, and increased mtCoQ reduction leading to increased H_2_O_2_ production during the oxidation of complex I (CI) and CII substrates. Statin treatment also altered the organization of the respiratory chain, leading to downregulation of the CI+CIII_2_+CIV and CIII_2_+CIV supercomplexes and decreased protein levels and activity of all respiratory chain complexes. Moreover, decreased cytochrome *a* + *a*_3_ content was accompanied by reduced maximum CIV activity. CoQ10 supplementation elevated mtCoQ levels, restored mitochondrial respiratory function, and decreased both mitochondrial H_2_O_2_ production and mtQ reduction level in the respiring mitochondria of statin-treated astrocytes. Altogether, the results show that prolonged statin exposure alters mtCoQ redox homeostasis and impairs mitochondrial bioenergetic function in astrocytes. Notably, CoQ10 supplementation attenuates these changes, supporting its potential role in protecting astrocyte mitochondria from statin-induced dysfunction.

## 1 Introduction

Statins are among the most commonly prescribed lipid-lowering drugs due to their proven effectiveness in reducing cardiovascular risk. By competitively inhibiting 3-hydroxy-3-methylglutaryl-coenzyme A reductase (HMG-CoA reductase) in the first step of the mevalonate pathway [1–3], statins reduce cholesterol biosynthesis and exert a broad spectrum of pleiotropic effects, including anti-inflammatory, antioxidant, and cytoprotective effects [4, 5]. Recent research indicates that statins may also affect the central nervous system (CNS), as lipophilic statins, such as atorvastatin and simvastatin, can cross the blood–brain barrier and alter CNS function [6].

Mitochondria are essential organelles that maintain cellular energy homeostasis. They generate ATP primarily through oxidative phosphorylation (OXPHOS), the final step of aerobic metabolism, driven by the mitochondrial respiratory chain. Coenzyme Q (CoQ) serves as a key electron transporter in the respiratory chain and is also an important lipid-soluble antioxidant present in cell membranes [7]. However, mitochondrial CoQ (mtCoQ) also participates in the production of mitochondrial reactive oxygen species (mtROS) at several mtCoQ-associated sites, particularly in complexes I (C1) and III (CIII) of the mitochondrial respiratory chain [8–10]. Alterations in mitochondrial activity, including decreased mtCoQ availability and impaired respiratory chain function, have been associated with cellular dysfunction and many metabolic, cardiovascular, and neurodegenerative diseases [11, 12].

Although statins are generally considered safe, increasing evidence suggests that they may affect mitochondrial metabolism [13, 14]. In addition to cholesterol production, the mevalonate pathway generates several important isoprenoid intermediates, including farnesyl pyrophosphate, geranylgeranyl pyrophosphate, heme *a*, and CoQ, which are essential for intracellular signaling, cell survival, and mitochondrial function [1, 3]. Statins’ inhibition of HMG-CoA reductase reduces the availability of these compounds and may thereby impair protein prenylation, mitochondrial maintenance, and OXPHOS [13, 14]. Moreover, the statin-induced decrease in mtCoQ levels may impair electron transport in the mitochondrial respiratory chain, reduce ATP synthesis, and promote excessive mtROS production, contributing to oxidative stress and mitochondrial dysfunction, as reported mainly in muscle cells [7, 13–18].

Astrocytes are the dominant glial cell population in the CNS and are essential for maintaining neuronal homeostasis, regulating neurotransmitter turnover, supporting synaptic activity, and providing metabolic substrates to neurons [19–22]. Although astrocytes primarily depend on aerobic glycolysis, mitochondrial OXPHOS is also important for supporting neuronal activity and maintaining brain homeostasis [23]. Therefore, changes in mitochondrial bioenergetics induced by statins’ inhibition of the mevalonate pathway may require metabolic adaptations in astrocyte mitochondria to maintain energy and redox homeostasis.

In addition to their lipid-lowering effects, statins exert multiple direct effects on astrocytes [24]. Lipophilic statins, such as atorvastatin and simvastatin, induce dose- and time-dependent stellation and apoptosis in cultured astrocytes, likely by eliminating isoprenoid intermediates [25]. Experimental studies have also shown that simvastatin reduces reactive astrogliosis, inhibits proinflammatory signaling, and suppresses neurotoxic astrocyte activation by modulating the nuclear factor kappa B (NF-κB) and Rho/Rho-associated coiled-coil containing protein kinase (ROCK) pathways [26–29]. These mechanisms may contribute to statins’ neuroprotective effects in CNS injury models [30].

Our recent study demonstrated that atorvastatin and simvastatin alter energy metabolism and redox balance in cultured astrocytes by reducing cellular CoQ content, modifying cellular respiration and mitochondrial turnover, lowering cellular ATP levels, and reducing ROS levels [31]. In addition, CoQ10 supplementation increases intracellular CoQ and ATP levels, supporting cellular energy metabolism. Although previous studies have shown that statins induce a significant deficiency in total intracellular CoQ levels in astrocytes, this study is the first to specifically examine statin-induced mtCoQ deficiency and its effects on mitochondrial bioenergetics, including mtCoQ redox status, mtROS production, and OXPHOS remodeling. Given astrocytes’ important role in brain energy homeostasis, understanding these mechanisms may provide further insights into statins’ effects on CNS function.

This study aimed to advance understanding of the mitochondrial effects of statin exposure in astrocytes by assessing mitochondrial respiratory parameters and functional activity and to better understand the potential neurobiological effects associated with long-term statin therapy. Building on our previous study at the cellular level [31], this study investigates the effects of long-term atorvastatin and simvastatin treatment on mitochondrial bioenergetics in cultured rat astrocytes, with particular focus on statin-induced mtCoQ deficiency. Using mitochondria isolated from control and statin-treated astrocytes, it assesses mitochondrial respiration, membrane potential (ΔΨ), OXPHOS efficiency, uncoupling, mtROS production, mtCoQ content and redox state, and the molecular organization of OXPHOS complexes to determine whether chronic statin exposure induces adaptive changes in mitochondria associated with decreased mtCoQ availability. Moreover, it investigates whether CoQ10 supplementation can restore mtCoQ levels and improve mitochondrial respiration and mtROS production in mitochondria of statin-treated astrocytes.

## 2 Experimental Procedures

### 2.1 Chemicals

All chemicals used in this study, including atorvastatin (catalog no. 1044516), simvastatin (catalog no. 1612700) and CoQ10 (C_59_H_90_O_4_; catalog no. 1705301), were obtained from Sigma-Aldrich (St. Louis, MO, USA). Detailed information on the antibodies is provided in Section 2.9. Stock solutions of statins and CoQ10 were prepared in methanol, which served as a vehicle. To obtain the active hydroxyacid form of simvastatin, the lactone ring was hydrolyzed in an alkaline medium, as previously described [31].

### 2.2 Astrocyte culture

The astrocyte cell line CTX TNA2 (catalog no. CRL-2006; ATCC, Manassas, VA, USA), derived from primary astrocytes isolated from the frontal cortex of 1-day-old rats, was used in this study. Cells were cultured under standard culture conditions for six days or treated with statins according to a previously described protocol [31]. Statin exposure consisted of 200 nM atorvastatin or 200 nM simvastatin. In experiments with CoQ10 supplementation, astrocytes were additionally treated with 3.4 µM CoQ10, where indicated. Under control conditions, cells were exposed to methanol (the vehicle) at an equivalent concentration (0.02%), without the addition of statins or CoQ10.

### 2.3 Mitochondria isolation

After 6 days of incubation, cells from the control and statin-treated groups were detached using a solution containing trypsin and ethylenediaminetetraacetic acid (EDTA). Next, the collected cells were washed sequentially with phosphate-buffered saline (PBS) containing 10% fetal bovine serum (FBS), then with 5% FBS in PBS, and finally with PBS alone, and then centrifuged at 1200 *g* and 4°C for 10 min. Then, they underwent three rounds of homogenization as follows: (i) cell pellet resuspension in PREP-I buffer (0.25 M sucrose, 1.5 mM EDTA, 1.5 mM ethylene glycol, bis[2-aminoethyl]tetraacetic acid [EGTA], 0.2% bovine serum albumin [BSA], 10 mM 4-[2-hydroxyethyl]-1-piperazineethanesulfonic acid [HEPES], and 5 mM Tris hydrochloride [Tris-HCl, pH 7.2]), (ii) disruption using a Teflon-glass homogenizer (18 strokes in round 1, 15 strokes in rounds 2 and 3), (iii) centrifugation at 1200 *g* for 10 min, and (iv) supernatant aspiration into a clean tube. Then, the supernatants containing the crude mitochondrial fraction were combined and centrifuged at 12,000 *g* for 10 min. Finally, the resulting mitochondrial pellets were washed with PREP-II buffer (0.25 M sucrose, 10 mM HEPES, and 5 mM Tris-HCl [pH 7.2]), recentrifuged at 12,000 *g* for 10 min, and resuspended in PREP-II buffer for further analysis.

### 2.4 Mitochondrial respiration and ΔΨ

Mitochondrial respiration and ΔΨ in isolated mitochondria were assessed according to previously published procedures [15, 32–34]. ΔΨ measurements were performed concurrently with oxygen consumption measurements using a tetraphenylphosphonium (TPP) selective electrode. Oxygen consumption rates (OCRs) were assessed polarographically using an oxygen electrode from Rank Bros. (Cambridge, UK) or Hansatech (King’s Lynn, UK). The measurements were performed at 37°C in a standard incubation medium (10 mM Tris-HCl [pH 7.2], 75 mM sucrose, 225 mM mannitol, 5 mM potassium chloride [KCl], 4 mM magnesium chloride [MgCl_2_], 5 mM monopotassium phosphate [KH_2_PO_4_], 0.15 mM EGTA, and 0.1% BSA). Depending on the oxygen electrode used, tests were performed in incubation volumes of 0.6 or 3.0 mL containing 0.6 or 3 mg of mitochondrial protein, respectively (final concentration of 1 mg/mL).

Phosphorylating respiration (state 3) was assessed in the presence of adenosine diphosphate (ADP) at 170 µM to determine the ADP/O ratio or 1.5 mM to induce maximal phosphorylating respiration. Uncoupled respiration was measured after the addition of carbonylcyanide-p-trifluoromethoxyphenylhydrazone (FCCP) at concentrations up to 1 µM. Resting non-phosphorylating respiration (state 4) was determined in the absence of ADP. Various substrate combinations were used to assess the activity of individual segments of the respiratory chain. CI-dependent respiration was measured using 3 mM malate + 3 mM pyruvate (CI substrates). Complex II (CII)-dependent respiration was measured using 3 mM succinate (CII substrate) combined with 1 µM rotenone (a CI inhibitor). CI- and CII-dependent respiration was jointly measured using 3 mM succinate, 3 mM malate, and 3 mM pyruvate. Maximal CIII activity was assessed using 0.3 mM duroquinol (+ 1 µM rotenone) in the presence of up to 1 µM FCCP.

Maximal complex IV (CIV; cytochrome *c* oxidase [COX]) activity was determined by monitoring the OCR using 0.15 mg of mitochondrial protein at a final concentration of 0.25 mg/mL. The assay was performed by sequential addition of specific reagents: including 4 µM antimycin A, 4 mM ascorbate, 0.06% cytochrome *c*, and increasing concentrations of *N*,*N*,*N*′,*N*′-tetramethyl-*p*-phenylenediamine (TMPD) up to 1.4 µM.

### 2.5 Energy-dissipating system activity

Uncoupling protein (UCP) and mitochondrial big-conductance Ca^2+^-regulated potassium channel (mitoBK_Ca_) activities were assessed by analyzing the relationship between oxygen consumption and ΔΨ during respiratory chain inhibition, as previously described [32]. To gradually reduce respiratory rate, oxidation of a mixture of malate, pyruvate, and succinate was inhibited by stepwise administration of rotenone (a CI inhibitor) and malonate (a CII inhibitor). The inhibitors were added sequentially in two steps: first, 0.25 µM rotenone with 0.8 mM malonate, then 0.5 µM rotenone with 1.6 mM malonate.

Non-phosphorylating respiration was measured in the presence of ATP synthase and ATP/ADP translocase inhibitors to minimize proton leakage associated with ATP turnover. Here, 0.8 µg/mL oligomycin and 0.8 µM carboxyatractyloside were added to the incubation medium. In UCP experiments, carboxyatractyloside was used to block fatty acid–induced proton transport by ATP/ADP translocase. UCP activity was assessed using 6 µM linoleic acid as an activator and 1.8 mM guanosine triphosphate (GTP) as an inhibitor. In addition, mitoBK_Ca_ activity was assessed using 0.8 µM NS11021 as an activator and 1.8 µM iberiotoxin (IbTx) as a selective inhibitor. UCP and mitoBK_Ca_ activities were quantified based on the flux–force relationship at the highest common ΔΨ value, representing linoleic acid–induced and GTP-sensitive UCP activity and NS11021-induced and IbTx-sensitive mitoBK_Ca_ activity, respectively.

### 2.6 mtCoQ concentrations and reduction levels

The concentrations of the mtCoQ species mtCoQ9 (the predominant form in rats) and mtCoQ10 (a much less abundant form), and the reduction levels of mtCoQ9 and mtQ10, were determined by high-performance liquid chromatography (HPLC), as previously described [35]. Separation was performed on a LiChrosorb RP-18 column (10 µm) using an ÄKTAexplorer fast protein liquid chromatography system (GE Healthcare, Chicago, IL, USA). Both oxidized and reduced forms of CoQ9 (mtCoQ9ox and mtCoQ9H_2_, respectively) and CoQ10 (mtCoQ10ox and mtCoQ10H_2_, respectively) were detected at 290 nm and 275 nm, respectively. Chromatographic peaks were identified and calibrated using commercial CoQ9 and CoQ10 standards. The reduction levels of mtCoQ9 and mtCoQ10 were calculated as the percentage of the reduced fraction (mtCoQH_2_) relative to the total mtCoQ pool (mtCoQtot). These measurements were performed during steady-state respiration with respiratory substrates. Samples used to mtCoQ extraction under conditions of respiratory chain activity (with respiratory substrates) were collected during measurements of oxygen consumption and ΔΨ. To determine the size of mtCoQH_2_ antioxidant pool (nonoxidizable by the respiratory chain) and the total amount of mtCoQ (mtCoQtot), measurements were performed under fully oxidizing conditions, in the absence of respiratory substrates, and thus with an inactive respiratory chain.

### 2.7 Mitochondrial H_2_O_2_ production

Hydrogen peroxide (H_2_O_2_) production in mitochondria isolated from astrocytes was quantified using the Amplex Red assay (Invitrogen, Waltham, MA, USA) in 24-well plates and a multimode microplate reader (Tecan Group Ltd., Männedorf, Switzerland). Measurements were performed at 37°C with 0.5 mg of mitochondrial protein (at a final concentration of 1 mg/mL) in 0.5 mL of standard incubation medium supplemented with 0.1 U/mL horseradish peroxidase (HRP), 4.5 µM Amplex Red, and 1 U/mL superoxide dismutase (SOD). Fluorescence signals were monitored for 40 min using excitation and emission wavelengths of 545 and 590 nm, respectively. Mitochondrial protein samples were incubated with various respiratory substrates: 3 mM malate + 3 mM pyruvate, 3 mM succinate + 1 µM rotenone, or 3 mM malate + 3 mM pyruvate + 3 mM succinate. Assays were performed in the presence or absence of 1.5 mM ADP to assess H_2_O_2_ production under phosphorylating and non-phosphorylating conditions, respectively. The mitochondrial H_2_O_2_ production rate was calculated via calibration with known H_2_O_2_ standards.

### 2.8 SOD activity

SOD2 activity was measured in mitochondria isolated from astrocytes using a commercial kit (CS0009, Sigma-Aldrich, St. Louis, MO, USA), in accordance with the manufacturer’s instructions. This method is based on the inhibition of the reduction of a tetrazolium salt to formazan – a process dependent on the superoxide anion – which is measured at a wavelength of 450 nm. Measurements were performed using 75 µg of mitochondrial protein in 96-well plates, with a multimode microplate reader (Tecan Group Ltd., Männedorf, Switzerland). SOD2 activity was expressed relative to control mitochondria isolated from astrocytes that had not been exposed to statins.

### 2.9 Mitochondrial cytochrome concentration and reduction

Mitochondria isolated from astrocytes were resuspended to a final protein concentration of 5 mg/mL in a buffer containing 120 mM KCl, 4 mM MgCl_2_, 20 mM Tris-HCl (pH 7.2), 2 µM rotenone, and 2.6 mM cyanide. The samples were then transferred to an experimental or reference cuvette. To determine the concentrations of individual cytochromes, complete reduction was achieved by adding 0.15% dithionite to the experimental cuvette. In the reference cuvette, complete oxidation was achieved by adding 1 mM potassium ferricyanide (K_3_[Fe(CN)_6_]). The level of cytochrome reduction was assessed by adding 0.3 mM duroquinol to the experimental cuvette to accelerate reduction via the cytochrome pathway. The difference spectra (reduced − oxidized) for cytochromes *c* + *c*_1_, *b*, and *a* + *a*_3_ were recorded using a UV spectrophotometer (model 1620; Shimadzu, Kyoto, Japan). The difference in absorbance was assessed between the following pairs of wavelengths: 552 and 540 nm (cytochrome *c* + *c*_1_), 563 and 577 nm (cytochrome *b*), and 605 and 630 nm (cytochromes *a* + *a*_3_). Difference spectra were used to determine both the cytochrome content and redox state.

### 2.10 Immunodetection of mitochondrial protein level

Proteins from mitochondrial fractions were resolved using 8%–12% sodium dodecyl sulfate–polyacrylamide gel electrophoresis (SDS-PAGE). Primary antibodies obtained from Abcam (Cambridge, UK) were used to immunodetect the following proteins: citrate synthase (CS, 46 kDa, catalog no. ab96600), coenzyme Q10B (COQ10B, 30 kDa, catalog no. ab41997), uncoupling protein 2 (UCP2, 31 kDa, catalog no. ab97931), CIII (ubiquinol-cytochrome c reductase core protein 2 [UQCRC2], 48 kDa, catalog no. ab203832), and total OXPHOS rodent WB antibody cocktail (catalog no. ab110413), which contains antibodies against subunits of CI (NADH:ubiquinone oxidoreductase subunit B8 [NDUFB8], 18 kDa), CII (succinate dehydrogenase complex iron sulfur subunit B [SDHB], 29 kDa), CIII (UQCRC2, 48 kDa), CIV (mitochondrially-encoded cytochrome *c* oxidase subunit I [MT-CO1/COX1], 38 kDa), and complex V (CV; ATP synthase peripheral stalk subunit F6 [ATP5PF/ATP5A], 54 kDa). A primary antibody obtained from Enzo Life Sciences (Farmingdale, NY, USA) was used to immunodetect superoxide dismutase 2 (SOD2, 24 kDa, catalog no. ADI-SOD). Primary antibodies obtained from Alomone Labs (Jerusalem, Israel) were used to immunodetect mitoBK_Ca_ channel subunits potassium calcium-activated channel subfamily M alpha 1 (KCNMA1/K_Ca_1.1, 105 kDa, catalog no. APC-107) and potassium calcium-activated channel subfamily M regulatory beta subunit 1 (KCNMB1/sloβ1, 37 kDa, catalog no. APC-036). Primary antibodies obtained from Thermo Fisher Scientific (Waltham, MA, USA) were used to immunodetect uncoupling protein 4 (UCP4, 53 kDa, catalog no. PA5-100668) and uncoupling protein 5 (UCP5, 37 kDa, catalog no. PA5-89394). CS was used as the loading control. Because a limited amount of mitochondrial material was available, the membranes were cut after transfer to allow separate immunodetection of target proteins and loading controls. The original uncropped immunoblots are provided in Supplementary Figs. S1–S5). Band intensities were quantified densitometrically using ImageJ software (National Institutes of Health, Bethesda, MD, USA).

### 2.11 BN-PAGE and in-gel activity assays

The in-gel activities of mitochondrial OXPHOS complexes CI, CII, CIV, and CIV were measured after separation of 100 µg of mitochondrial proteins by blue native polyacrylamide gel electrophoresis (BN-PAGE) with a gel gradient of 3.0%–9.5%, according to previously established procedures [15]. The BN-PAGE-separated proteins were also transferred onto nitrocellulose membranes to determine levels of CIII supercomplexes by immunoblotting with an anti-UQCRC2 antibody (catalog no. ab14745; Abcam, Cambridge, UK).

### 2.12 Statistical analysis

All statistical analyses were performed using SPSS Statistics (IBM Corp., Armonk, NY, USA). Data are presented as means ± standard deviations (SD), obtained from at least 5–6 independent biological experiments. Each experiment involved mitochondria isolated from a separately cultured and treated population of CTX TNA2 astrocytes. All measurements were performed in at least three technical replicates, which were averaged to obtain a single value for each biological replicate. Thus, independent biological experiments (n = 5–6), rather than technical replicates, were considered the experimental units for statistical analysis. Data normality was assessed using Q–Q plots and the Shapiro–Wilk test. Differences among experimental groups were evaluated using one-way analysis of variance (ANOVA), followed by Tukey’s post hoc test for multiple comparisons. Differences were considered statistically significant at *P* < 0.05. Statistical significance is indicated as follows: *, *P* < 0.05; **, *P* < 0.01; ***, *P* < 0.001 compared with control mitochondria; ^#^, *P* < 0.05, ^##^, *P* < 0.01, ^###^, *P* < 0.001 for comparison between the indicated groups of mitochondria.

## 3 Results

Lipophilic statins are better able to cross the blood–brain barrier than hydrophilic statins, allowing them to affect the cells in the CNS, including astrocytes [36–39]. Therefore, two commonly prescribed lipophilic statins were selected for this study: atorvastatin, administered in its pharmacologically active hydroxyacid form, and simvastatin, which is administered as a lactone prodrug and undergoes first-pass hepatic hydrolysis to form the active β-hydroxyacid metabolite [40]. Because simvastatin must be converted to the active form before use in cell culture experiments, it was chemically activated before in vitro administration. The nanomolar concentrations used in the experiments were chosen based on previously published reports describing serum concentrations achieved in patients undergoing standard therapeutic statin treatment [41–44]. All experiments were performed using astrocytes cultured for 6 days under control conditions or in the presence of 200 nM atorvastatin or simvastatin. We have previously shown that these concentrations do not affect astrocyte viability while inducing a comparable significant reduction in cellular CoQ levels (∼30% in CoQ9 content and ∼70% in CoQ10 content) [31]; higher statin concentrations (≥250 nM) resulted in pronounced cytotoxic effects, significantly reducing astrocyte viability and markedly decreasing total cellular CoQ levels (∼60%).

### 3.1 Statins decrease the levels of mtCoQtot and mtCoQH_2_, but do not affect the level or activity of SOD2 in astrocyte mitochondria

The effects of statins on mtCoQ homeostasis in astrocytes remain poorly understood. To investigate whether prolonged statin exposure alters mtCoQ content, we quantified mtCoQ9 and mtCoQ10 levels in mitochondria isolated from astrocytes cultured for 6 days in the presence of 200 nM atorvastatin or simvastatin. Both statins significantly reduced mtCoQtot levels compared with untreated controls. Specifically, total mtCoQ9 (mtCoQ9tot) content (mtCoQ9H_2_ + mtCoQ9ox) decreased by ∼30%, and total mtCoQ10 (mtCoQ10tot) content (mtCoQ10H_2_ + mtCoQ10ox) decreased by ∼35% (Fig. 1a).

**FIGURE 1.**
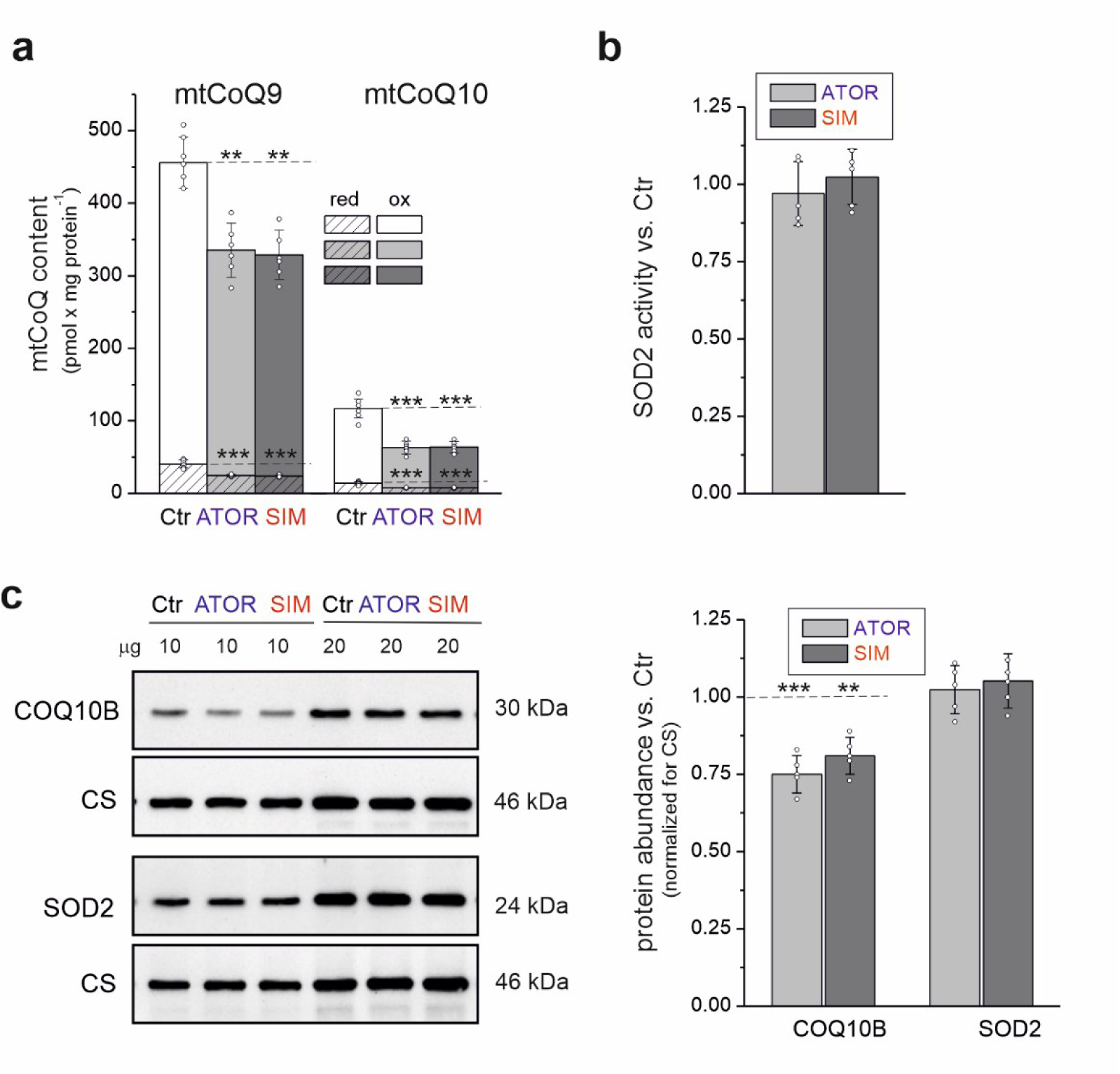
(**a**) mtCoQ9 and mtCoQ10 contents, SOD2 activity, and (**c**) COQ10B and SOD2 levels in mitochondria isolated from astrocytes treated for 6 days with 200 nM atorvastatin (ATOR) or simvastatin (SIM), compared with those from untreated control astrocytes (Ctr). **a**, Reduced (mtCoQH_2_), oxidized (mtCoQox), and total (mtCoQH_2_ + mtCoQox) CoQ contents were determined under fully oxidizing conditions in the absence of respiratory substrates capable of reducing mtCoQ. **b**, SOD2 activity expressed relative to control mitochondria. **c**, Quantification of protein levels and representative immunoblot images. Loading control: CS. Full, uncropped immunoblots are provided in Supplementary Fig. S1. **a**-**c**, Data are presented as the mean ± SD (*n* = 5–6). Significance: **, *P* < 0.01; ***, *P* < 0.001 compared with Ctr (horizontal lines).

To further assess changes in mitochondrial redox status, we analyzed mtCoQH_2_ content under fully oxidizing conditions in the absence of respiratory substrates that could reduce mtCoQ. This fraction represents mtCoQ, which is resistant to oxidation by the respiratory chain and may constitute an antioxidant pool [7]. In mitochondria isolated from untreated control astrocytes, mtCoQ9H_2_ content accounted for ∼9% of mtCoQ9tot, and mtCoQ10H_2_ accounted for ∼12% of mtCoQ10tot. Notably, these contents were ∼40% lower in mitochondria obtained from atorvastatin- or simvastatin-treated astrocytes (Fig. 1a), indicating a significant reduction in the antioxidant fraction of mtCoQ after statin exposure. Given the important role of CoQ-related proteins in maintaining respiratory chain function, we next examined the levels of COQ10B, a protein that supports mtCoQ function. COQ10B levels were significantly lower (∼20%) in mitochondria from statin-treated astrocytes than in those from untreated control astrocytes (Fig. 1b).

Because a significant decrease in mtCoQH_2_ content could increase oxidative stress, we also assessed mitochondrial activity and the levels of the antioxidant enzyme SOD2. Despite decreased mtCoQ9H_2_ and mtCoQ10H_2_ contents, both the activity and protein levels of SOD2 remained unchanged after statin treatment (Fig. 1b, c). Thus, although statins significantly impaired mtCoQ availability and mitochondrial redox status, they did not induce compensatory changes in the levels or activity of the mitochondrial SOD2 under the examined experimental conditions.

Because mtCoQH₂ acts not only as a major mitochondrial antioxidant but also as an essential electron carrier in the respiratory chain, we next investigated whether statin-induced decreases in mtCoQ content affect mitochondrial bioenergetic function, including mitochondrial respiration, ATP synthesis efficiency, and ΔΨ.

### 3.2 Statins decrease respiratory activity, ΔΨ, and ATP synthesis during CI- and CII-dependent respiration in astrocyte mitochondria

Mitochondrial respiration was assessed during the oxidation of malate and pyruvate (CI substrates); succinate (CII substrate), in the presence of rotenone (a CI inhibitor); and malate, pyruvate, and succinate under both non-phosphorylating and phosphorylating conditions. Analysis of mitochondrial coupling parameters revealed that chronic exposure of astrocytes to atorvastatin or simvastatin altered mitochondrial respiratory activity under all examined substrate conditions (Table 1). Although the ADP/O ratio remained unchanged, the respiratory control ratio (RCR) was ∼8%–16% lower in mitochondria from statin-treated astrocytes. Moreover, ADP phosphorylation was lower (20%–25%) in mitochondria isolated from statin-treated astrocytes than in those isolated from untreated control astrocytes, regardless of the substrates (Table 1). These results demonstrate that long-term exposure to atorvastatin or simvastatin significantly impairs mitochondrial ATP production. Under non-phosphorylating conditions, mitochondria from statin-treated astrocytes showed a slower respiratory rate (∼12%) only during the oxidation of malate and pyruvate, substrates associated with CI, compared with those from untreated control astrocytes (Fig. 2a); they showed unaltered respiratory rates during the oxidation of succinate or mixed substrates. However, the effects of statin treatment became more pronounced under phosphorylating conditions. Respiratory rates were significantly lower (∼15%–20%) in mitochondria from statin-treated astrocytes than in those from untreated control astrocytes for all examined substrate combinations. Under both non-phosphorylating and phosphorylating conditions, ΔΨ was significantly lower in mitochondria from statin-treated astrocytes than in those from untreated control astrocytes for all examined substrate combinations (Fig. 2b).

**FIGURE 2.**
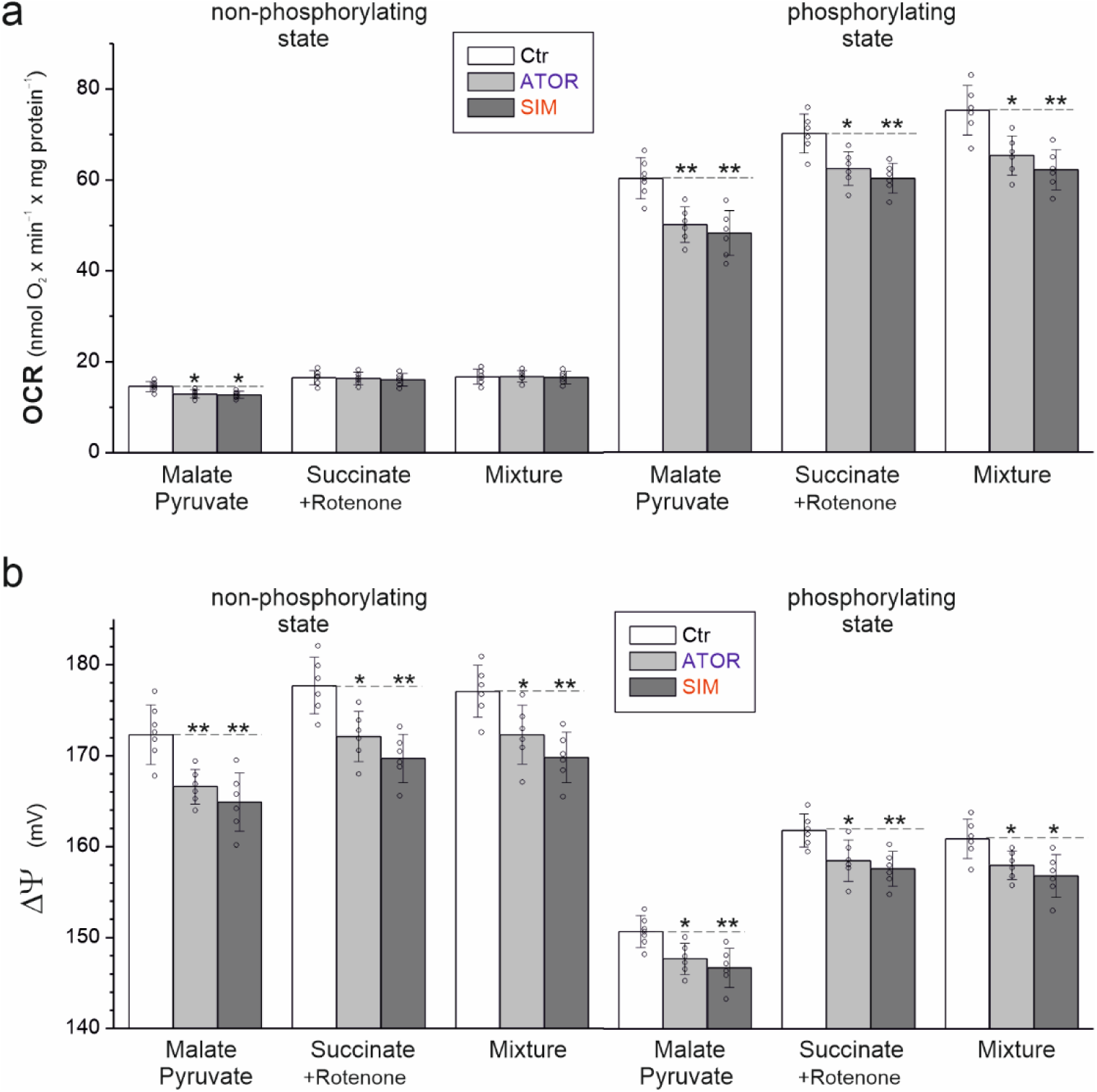
(**a**) OCR and (**b**) ΔΨ of mitochondria isolated from untreated control astrocytes (Ctr) and atorvastatin (ATOR)- or simvastatin (SIM)-treated astrocytes during the oxidation of various respiratory substrates under non-phosphorylating and phosphorylating conditions. **a**, **b**, Data are presented as the mean ± SD (*n* = 6). Significance: *, *P* <0.05; **, *P* <0.01 compared with Ctr (horizontal lines).

**Table 1.** Mitochondrial bioenergetic coupling parameters, including respiratory control ratio (RCR), ADP/O ratio, and ADP phosphorylation rate (calculated as phosphorylating respiration × ADP/O), measured in mitochondria isolated from control astrocytes (Ctr) and astrocytes treated with atorvastatin (ATOR) or simvastatin (SIM). ADP phosphorylation rate is presented as nmol ADP × min^−1^ × mg protein^−1^. Mean ± SD (*n* = 6); statistics: one-way ANOVA, \**P* < 0.05, \*\**P* < 0.01, *** *P* < 0.001 compared with control mitochondria.

|  | **Malate + Pyruvate** |  |  | **Succinate + Rotenone** |  |  | **Malate + Pyruvate + Succinate** |  |  |
|---|---|---|---|---|---|---|---|---|---|
|  | Ctr | ATOR | SIM | Ctr | ATOR | SIM | Ctr | ATOR | SIM |
| **RCR** | 4.12 ± 0.03 | 3.86 ± 0.03<br>*** | 3.77 ± 0.16<br>** | 4.25 ± 0.15 | 3.82 ± 0.10<br>*** | 3.75 ± 0.13<br>*** | 4.49 ± 0.13 | 3.89 ± 0.04<br>*** | 3.76 ± 0.05<br>*** |
| **ADP/O** | 2.38 ± 0.19 | 2.31 ± 0.13 | 2.28 ± 0.17 | 1.31 ± 0.07 | 1.25 ± 0.07 | 1.21 ± 0.06 | 1.99 ± 0.11 | 1.91 ± 0.11 | 1.85 ± 0.09 |
| **Phosphorylation rate** | 144 ± 23 | 116 ± 16<br>* | 111 ± 19<br>* | 91.7 ± 11 | 78.1 ± 9.2<br>* | 73.0 ± 7.8<br>** | 150 ± 20 | 125 ± 16<br>* | 115 ± 14<br>** |

Statin treatment led to a decrease in the respiration rate under phosphorylating conditions, the rate of ADP phosphorylation, and RCR, while having no significant effect on the ADP/O ratio (Table 1). These findings indicate that statins reduce mitochondrial OXPHOS capacity, while maintaining the stoichiometric efficiency of ATP synthesis. A reduced RCR primarily reflects an impairment in ADP-stimulated respiration, whereas respiration under non-phosphorylating conditions remains unchanged during oxidation of CII and combined CI+CII substrates. Thus, statin-induced mitochondrial dysfunction appears to affect the capacity of the OXPHOS system rather than the phosphorylation yield, indicating that mitochondria produce ATP at a lower rate without a proportional decline in ATP synthesis efficiency.

Altogether, these results indicate that mitochondria isolated from astrocytes chronically exposed to atorvastatin or simvastatin exhibit lower respiratory activity, reduced ΔΨ, and decreased ATP synthesis compared with those from untreated control astrocytes. They are consistent with CoQ serving as a central electron transporter, linking CI and CII to downstream components of the mitochondrial respiratory chain, and indicate that decreased CoQ availability impairs both CI- and CII-dependent mitochondrial respiration.

Because mtCoQH_2_ not only acts as an important antioxidant but also participates in the production of mtROS through electron transport in the mitochondrial respiratory chain, changes in its redox state may influence mtROS production. Therefore, we next investigated whether the statin-induced decrease in mtCoQ in astrocytes affects the level of mtCoQ reduction and thereby mtROS production.

### 3.3 Statins increase mtROS production and mtCoQ reduction level in astrocyte mitochondria

Mitochondria isolated from astrocytes chronically exposed to atorvastatin or simvastatin showed consistently elevated H_2_O_2_ production compared with mitochondria from control cells (Fig. 3a). This increase was observed under both non-phosphorylating and phosphorylating conditions and occurred regardless of the respiratory substrate used, including malate and pyruvate (CI substrates), succinate (CII substrate), or a combination of CI and CII substrates. However, the magnitude of this effect depended on the oxidized substrate. Comparisons between individual substrate conditions showed that the increase in H_2_O_2_ production was more pronounced during the oxidation of CI-related substrates (an ∼30–40% increase), whereas the oxidation of CII substrate, resulted in a smaller increase of ∼12–18%.

**FIGURE 3.**
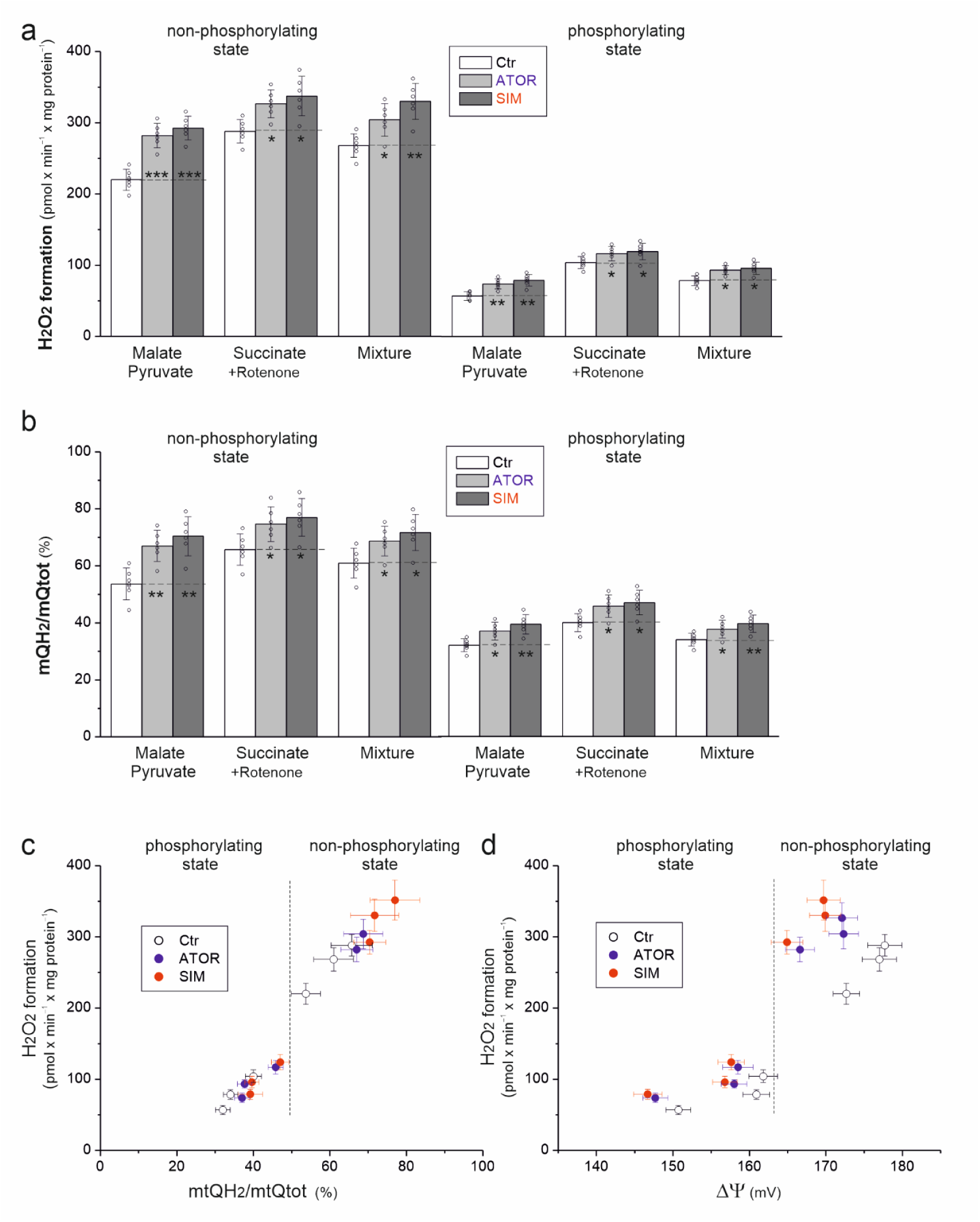
(**a**) H_2_O_2_ production rate, (**b**) mtCoQ reduction level (mtCoQH_2_/mtCoQtot), (**c**) relationships between H_2_O_2_ production and the mtCoQ reduction level and H_2_O_2_ production, and (**d**) ΔΨ of mitochondria isolated from untreated control astrocytes (Ctr) and atorvastatin (ATOR)- or simvastatin (SIM)-treated astrocytes during the oxidation of various respiratory substrates under non-phosphorylating and phosphorylating conditions. **a**-**d**, Data are presented as the mean ± SD (*n* = 6). Significance (**a**, **b**): *, *P* <0.05; **, *P* <0.01; ***, *P* <0.001 compared with Ctrl (horizontal lines).

Mitochondria isolated from astrocytes chronically exposed to atorvastatin or simvastatin showed consistently elevated H_2_O_2_ production compared with mitochondria from untreated control astrocytes (Fig. 3a). This increase was observed under both non-phosphorylating and phosphorylating conditions and occurred regardless of the respiratory substrate used (malate and pyruvate, succinate, or their combination). However, the magnitude of this effect depended on the oxidized substrate. Comparisons between individual substrate conditions showed that the increase in mitochondrial H_2_O_2_ production was greater during the oxidation of CI-related substrates (∼30%–40%) compared with the CII substrate (∼12%–15%). To elucidate the mechanism underlying the increased production of mtROS in the mitochondria of astrocytes treated with statins, we also measured H_2_O_2_ release during succinate oxidation in the absence of rotenone (Supplementary Fig. S2b). Under both non-phosphorylating and phosphorylating conditions, the greater increase in H_2_O_2_ release observed in the absence of CI inhibitor (25–27%), compared to the increase recorded in its presence (12–15%), indicates the involvement of reverse electron transport (RET) in mtROS production.

To further support the mitochondrial origin of the ROS signal measured using the Amplex Red reagent (Fig. 3a), we conducted additional experiments using MitoSOX Red, a mitochondria-targeted fluorescent probe designed to assess mitochondrial superoxide anion production (Supplementary Fig. 2Sb). The results obtained were consistent with those from the analysis of H_2_O_2_ production using the Amplex Red reagent, providing further evidence of increased ROS production in mitochondria isolated from astrocytes treated with statins compared to control mitochondria.

Moreover, mitochondria from statin-treated astrocytes consistently showed a greater level of mtCoQ reduction (mtCoQH_2_/mtCoQtot) than mitochondria from untreated control astrocytes under both respiratory states and all examined substrate conditions (Fig. 3b). Like for mtROS, substrate-dependent differences were observed under non-phosphorylating conditions. The increase in mtCoQ reduction was greatest during the oxidation of CI substrates (∼25%–30%) compared with the CII substrate (∼12%–17%). Moreover, the CoQH_2_/CoQox ratio was also higher in mitochondria from statin-treated astrocytes than in mitochondria from untreated control astrocytes, regardless of the respiratory state and for all substrates tested (Supplementary Fig. S3). This increase was due to the fact that the decline in CoQH_2_ content was smaller than the decline in CoQox content.

These results indicate that chronic exposure of astrocytes to statins increases mtCoQ reduction and mtROS production. The stronger effects observed during CI-dependent respiration suggest that statin-induced disruption of mtCoQ homeostasis preferentially impairs CI-dependent electron flow, thereby increasing mtROS production.

Under all examined substrate oxidation conditions, mitochondria isolated from untreated control and statin-treated astrocytes showed a single relationship between H_2_O_2_ production and the mtCoQ reduction level, expressed as either mtCoQred/mtCoQtot or mtCoQH_2_/mtCoQox, under both non-phosphorylating and phosphorylating conditions (Fig. 3c, Supplementary Fig. S3d). This observation indicates that mtROS production is directly dependent on the level of mtCoQ reduction [45]. However, analysis of the relationship between H_2_O_2_ production and ΔΨ (Fig. 3d) revealed a marked upward shift in H_2_O_2_ production but downward shift in ΔΨ in mitochondria from statin-treated astrocytes compared with those from untreated control astrocytes (Fig. 3d). This pattern indicates that inhibition of the mtCoQH_2_-oxidizing cytochrome pathway (CIII and CIV) leads to increased mtROS production.

### 3.4 Statins decrease cytochrome pathway activity and cytochrome *a* + a_3_ content in astrocyte mitochondria

To better characterize the mechanism underlying impaired mitochondrial function in statin-treated astrocytes, we examined CIII and CIV activity. Mitochondria isolated from atorvastatin- or simvastatin-treated astrocytes showed significantly lower CIII and CIV activity (∼16%–18%) than those isolated from untreated control astrocytes (Fig. 4a). We also assessed the level of reduction of respiratory chain cytochromes using duroquinol as an artificial electron donor. Mitochondria from statin-treated astrocytes showed significantly less reduction of all analyzed cytochromes, including cytochromes *c* + *c*_1_, *b*, and *a* + *a*_3_ (Fig. 4c), supporting the observed reduction in CIII and CIV activity.

**FIGURE 4.**
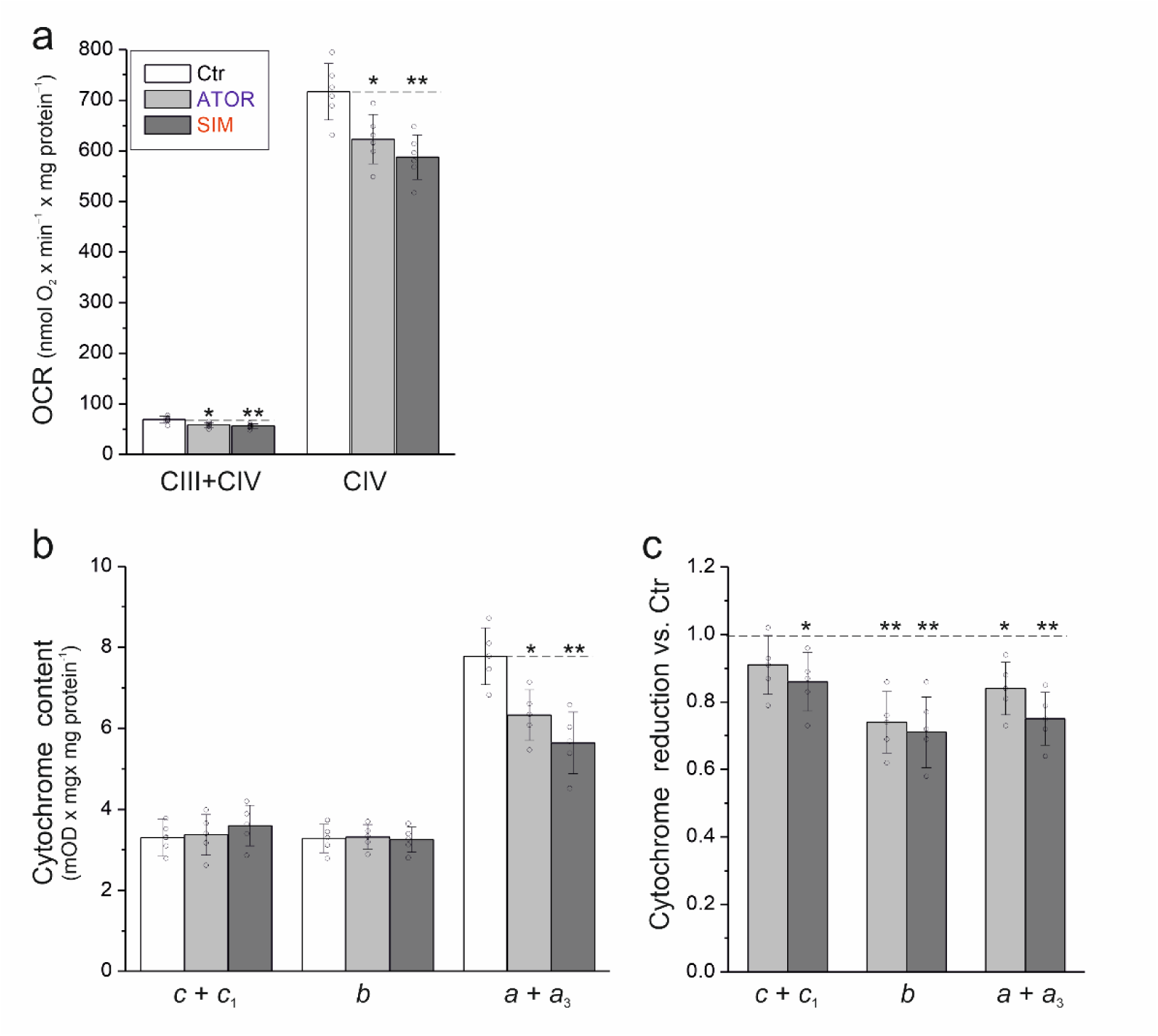
(**a**) Maximal CIII and CIV activities (OCR), (**b**) cytochrome content, and (**c**) cytochrome reduction in mitochondria isolated from untreated control astrocytes (Ctr) and atorvastatin (ATOR)- or simvastatin (SIM)-treated astrocytes. **a–c**, Data are presented as the mean ± SD (*n* = 5–6). Significance: *, *P* <0.05; **, *P* <0.01 compared with Ctr (horizontal lines).

Because statins inhibit the mevalonate pathway and reduce the synthesis of metabolites other than CoQ, including isoprenoid intermediates essential for heme *a* biosynthesis, we investigated whether they affect other components of the mitochondrial respiratory chain. Because heme *a* is an essential prosthetic group of CIV, we analyzed mitochondrial cytochrome content. While cytochrome *b* and *c* + *c*_1_ content remained unchanged, cytochrome *a* + *a*_3_ content was ∼20% lower in the mitochondria from statin-treated astrocytes compared with those from untreated control astrocytes (Fig. 4b).

Altogether, these results suggest that statin-induced inhibition of the mevalonate pathway may affect not only CoQ biosynthesis but also reduce the availability of heme *a* in mitochondria. Consequently, statin-induced impairment of mitochondrial respiration may result from deficiencies in mtCoQ and heme *a*, contributing to reduced CIV activity and impaired electron transport chain function in astrocytes.

### 3.5 Statins alter the molecular organization of OXPHOS in astrocyte mitochondria, downregulating the respiratory chain CI+CIII_2_+CIV and CIII_2_+CIV supercomplexes and CII

To further investigate statin-induced changes in mitochondrial bioenergetics, we examined the expression and functional activity of individual OXPHOS components and their supramolecular organization. Immunoblot analysis revealed an ∼10%–18% decrease in the protein levels of CI (NDUFB8 subunit), CII (SDHB subunit), CIII (Core 2 subunit; only for mitochondria from simvastatin-treated astrocytes), and CIV (COXI subunit) in mitochondria isolated from statin-treated astrocytes (Fig. 5a). In contrast, the protein levels of CV (α subunit) remained unchanged. To assess the functional consequences of these changes, we performed BN-PAGE and in-gel activity assays. The activities of the CI+CIII_2_+CIV and CIII_2_+CIV supercomplexes, as well as CII, were significantly lower in mitochondria from atorvastatin- or simvastatin-treated astrocytes (Fig. 5b). In contrast, the activities of CI, CIII, and CIV outside the supercomplex assemblies, as well as CV, were preserved.

**FIGURE 5.**
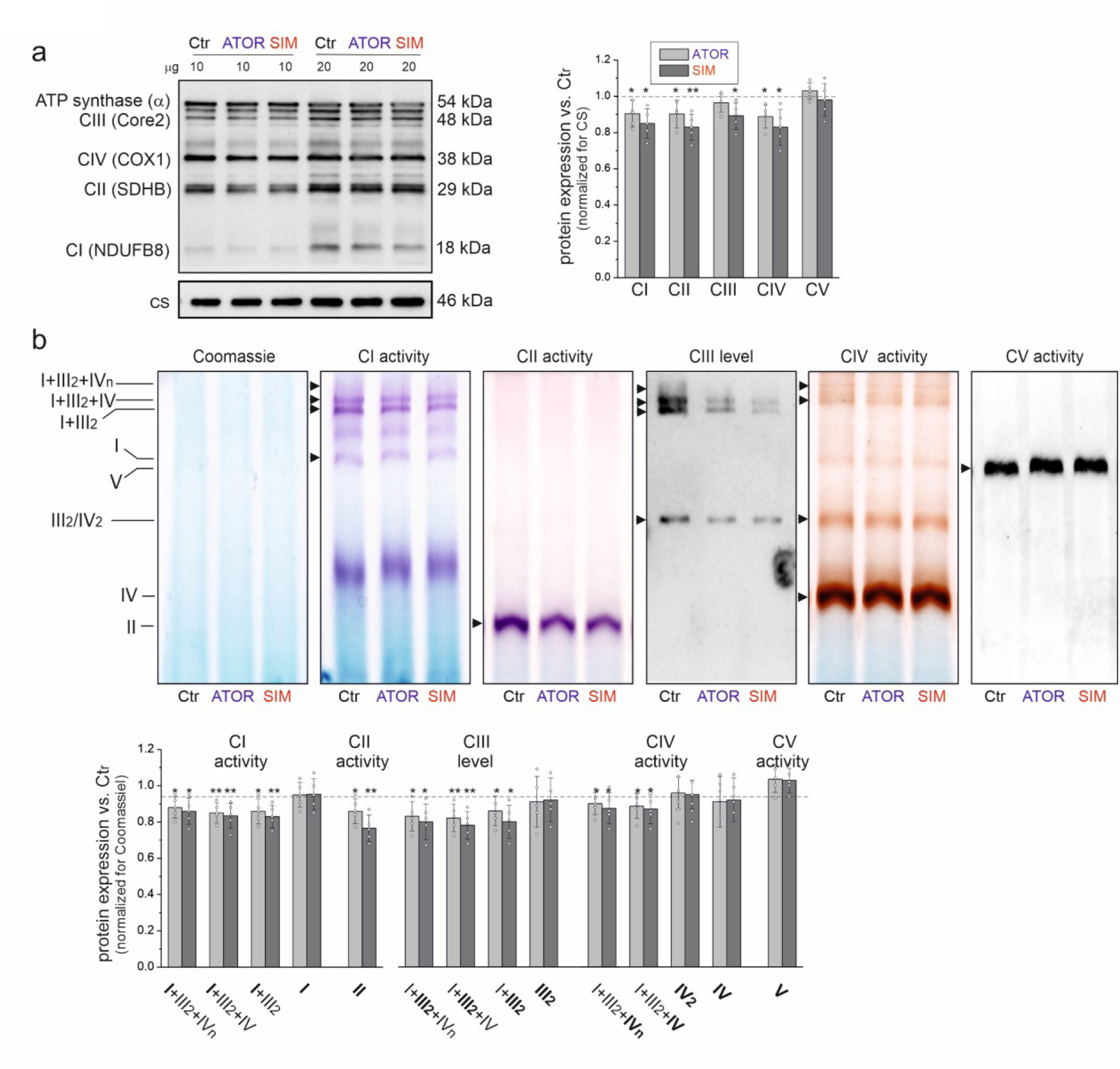
(**a**) Complexes of the OXPHOS system and (**b**) supercomplexes/complexes in the respiratory chain of mitochondria isolated from untreated control astrocytes (Ctr) and atorvastatin (ATOR)- or simvastatin (SIM)-exposed astrocytes. **a**, Representative immunoblot and protein abundance analysis. **b**, Representative BN-PAGE showing the supercomplexes and complexes of the OXPHOS system (upper part) and changes in in-gel activity (CI, CII, CIV, and CV) or protein levels (CIII) (lower part). Data are presented as the mean ± SD (*n* = 5). Significance: *, *P* < 0.05; **, *P* <0.01 compared with Ctr. The original immunoblots and gels are shown in Supplementary Fig. S4. CI–CIV, respiratory chain complexes; CV, ATP synthase.

These results are consistent with reduced CIII (Core 2 subunit) abundance observed by SDS-PAGE (Fig. 5a) and reduced CIII enzymatic activity measured during duroquinol oxidation in intact mitochondria (Fig. 4a). Moreover, the lower activity of CIV-associated supercomplexes in the gel correlated with decreased COXI protein levels (Fig. 5), decreased CIV activity measured in intact mitochondria (Fig. 4a), and decreased cytochrome *a* + *a_3_* content/reduction (Fig. 4b, c).

Altogether, these results demonstrate that prolonged statin exposure, which significantly lowers mtCoQ levels in astrocyte mitochondria, induces significant remodeling of respiratory chain supercomplex organization. Notably, statin-induced decreases in mtCoQ downregulate the CI+CIII_2_+CIV and CIII_2_+CIV supercomplexes and CII, thereby contributing to mitochondrial dysfunction in astrocytes. These changes may represent an adaptive response aimed at limiting mtROS production at mtCoQ-dependent sites in CI and CIII.

### 3.6 Statins do not affect energy-dissipating systems in astrocyte mitochondria

Because mitochondria from astrocytes treated with atorvastatin and simvastatin showed increased mtROS production (Fig. 3a), we next investigated whether statin exposure alters the activity or expression of two mitochondrial energy-dissipating systems: UCPs and mitoBK_Ca_ channel. These systems have been proposed to exert their antioxidant effects by mildly dissipating proton force and thereby increasing electron flow in the respiratory chain [32, 46, 47]. Functional analysis using the flux‒force kinetics assays showed that neither fatty acid-induced, GTP-sensitive UCP activity nor NS11021-stimulated, IbTx-sensitive mitoBK_Ca_ activity differed between mitochondria isolated from statin-treated and untreated control astrocytes (Fig. 6a, b). Consistent with the functional data, immunoblot analysis revealed no significant differences in UCP2, UCP4, or UCP5 protein abundance between mitochondria from statin-treated and untreated control astrocytes (Fig. 6c). Similarly, statin treatment did not affect the expression of the pore-forming α-subunit (K_Ca_1.1) and regulatory subunit (sloβ1) of mitoBK_Ca_.

**FIGURE 6.**
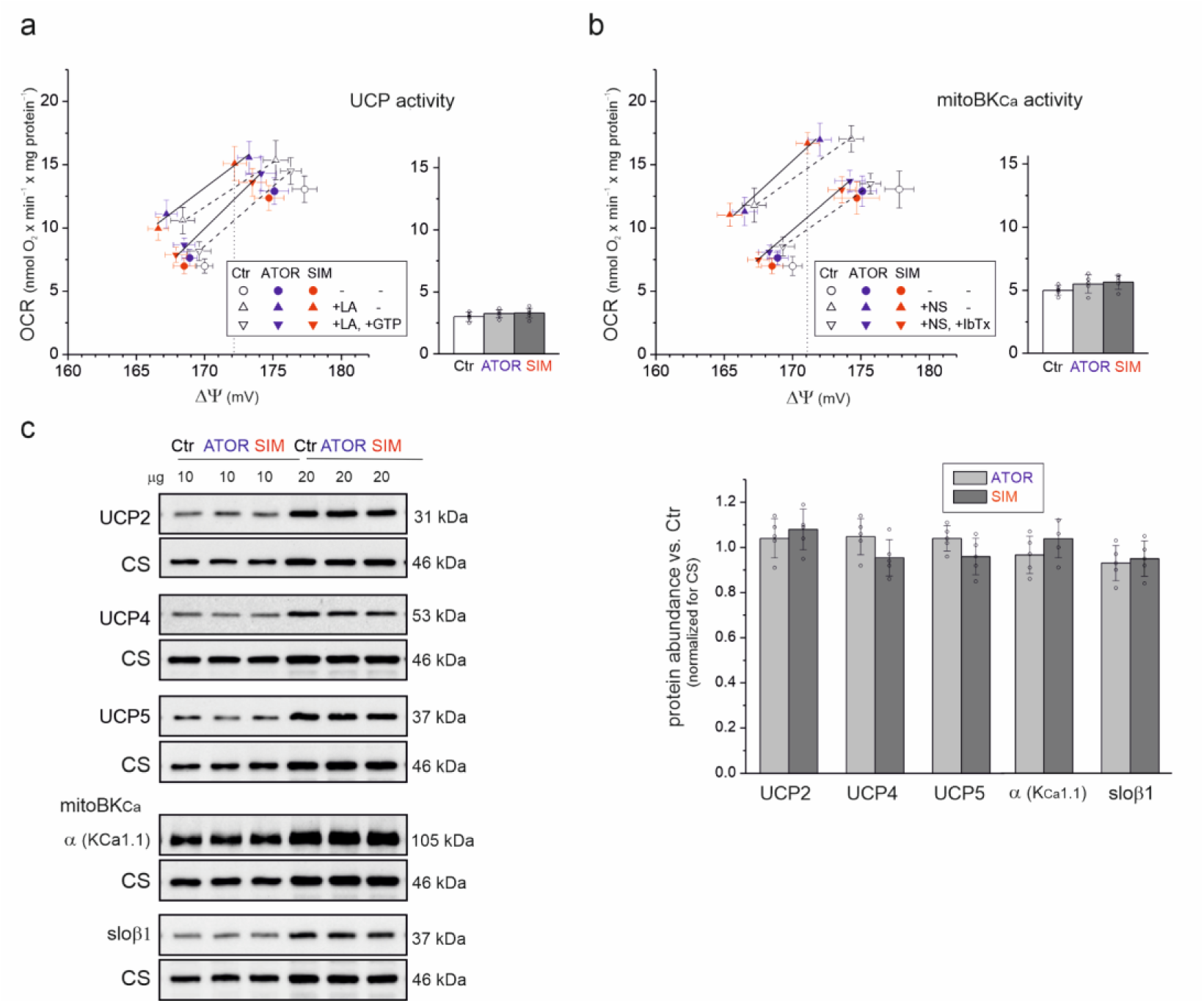
(**a**, **b**) Activities and (**b**) protein levels of energy-dissipating systems UCP and mitoBK_Ca_ in the mitochondria of untreated control astrocytes (Ctr) and atorvastatin (ATOR)- or simvastatin (SIM)- treated astrocytes. **a**, **b**, Relationships between the OCR and ΔΨ for malonate + rotenone titration during the oxidation of a mixture of malate and pyruvate, and under non-phosphorylating conditions. UCP and mitoBK_Ca_ activity were determined at the same ΔΨ (∼172 mV). **a**, UCP activity was calculated as linoleic acid-induced GTP-inhibited UCP-mediated H^+^ leakage. **b**, mitoBK_Ca_ activity was calculated as NS11021-induced IbTx-inhibited OCR. **c**, Representative immunoblots and analyses of protein abundance. Subunits of mitoBK_Ca_: pore-forming α subunit (K_Ca_1.1) and auxiliary subunit (sloβ1). Full, uncropped immunoblots are provided in Supplementary Fig. S5. Data are presented as mean ± SD (*n* = 5–6).

Altogether, the functional and protein expression findings indicate that chronic exposure of astrocytes to atorvastatin or simvastatin does not activate mitochondrial energy dissipation systems. Thus, despite increased mtROS production, these systems are not involved in the mitochondrial response to statin treatment.

### 3.7 CoQ10 supplementation elevates CoQ levels and restores respiration and mtROS production in statin-treated astrocyte mitochondria CoQ10 supplementation elevates

We next examined how CoQ10 supplementation affects the redox states of mtCoQ9 and mtCoQ10 in the mitochondria of statin-treated astrocytes (Fig. 7a, b). Because previous experiments revealed no significant differences between atorvastatin and simvastatin treatment (Figs. 1–6), only simvastatin was examined in the CoQ10 supplementation experiments. Mitochondria were isolated from astrocytes cultured in media with or without 3.4 µM CoQ10, a concentration comparable to the plasma concentration in human patients receiving CoQ10 supplementation [48].

**FIGURE 7.**
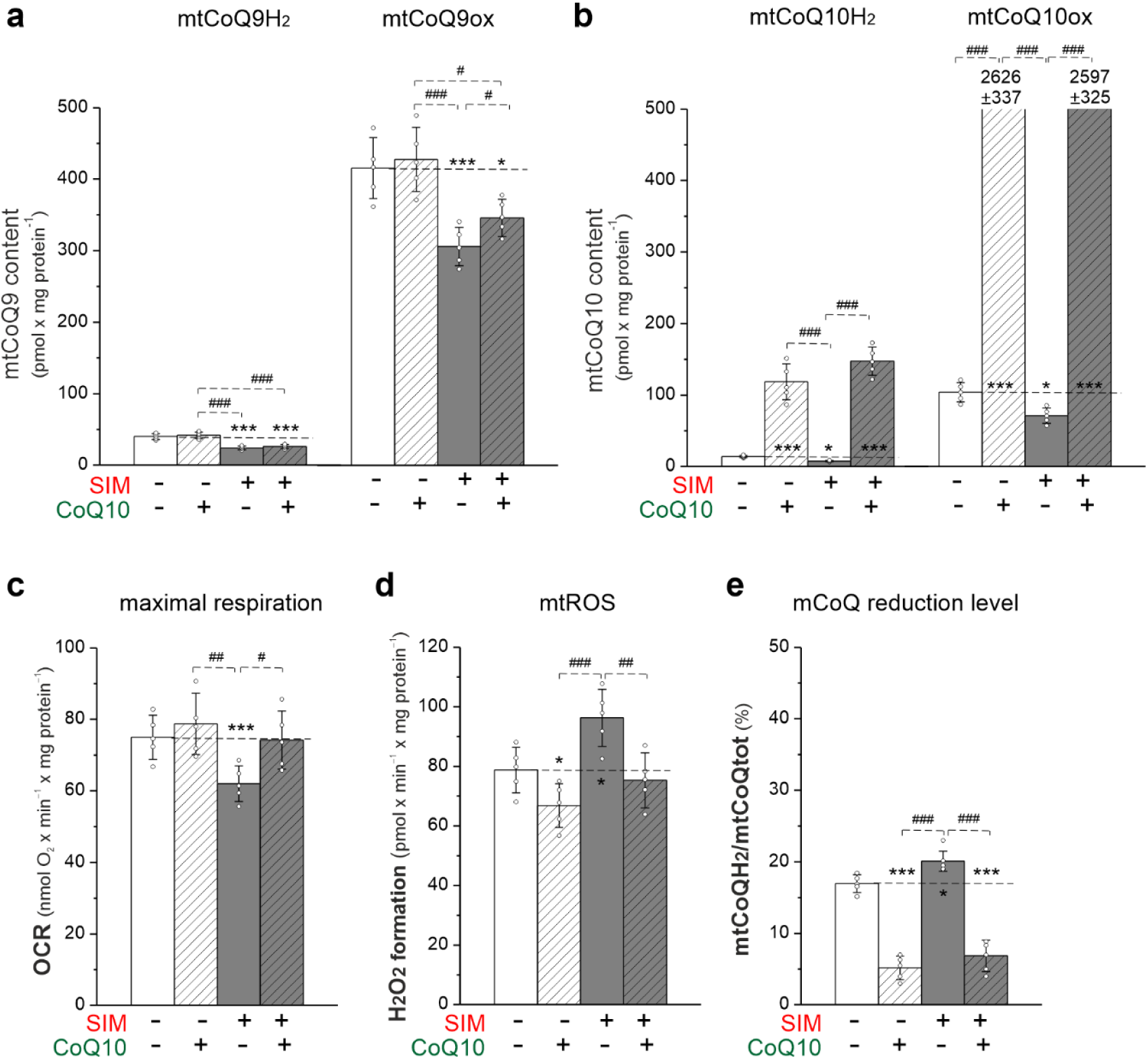
Effects of CoQ10 supplementation on (**a**) mtCoQ9 and (**b**) mtCoQ10 content, (**c**) maximal respiration, (**d**) mtROS production, and (**e**) mtCoQ reduction level in mitochondria from control (Ctr), simvastatin (SIM)-treated, and CoQ10+SIM-treated astrocytes. Mitochondria were isolated from astrocytes cultured for 6 days with or without 200 nm SIM and/or 3.4 µM CoQ10. **a**, **b**, Content of reduced (mtCoQH_2_) and oxidized (mtCoQox) forms of mtCoQ9 and mtCoQ10 were determined under fully oxidizing conditions in the absence of respiratory substrates capable of reducing mtCoQ. **c**-**e**, OCR, H_2_O_2_ production, and mtCoQ reduction level (mtCoQH_2_/mtCoQtot) were measured with a mixture of respiratory substrates (i.e., 3 mM malate, 3 mM pyruvate and 3 mM succinate) under phosphorylating conditions (i.e., in the presence of 1.5 mM ADP). **a–e**, Data are presented as the mean ± SD (*n* = 5). Significance: *, *P* < 0.05; **, *P* < 0.01; ***, *P* < 0.001 compared with Ctr (horizontal lines); ^#^, *P* < 0.05, ^##^, *P* < 0.01, ^###^, *P* < 0.001 between indicated groups.

To determine the size of mtCoQH_2_ antioxidant pool (nonoxidizable by the respiratory chain) and the total amount of mtCoQ, measurements were performed under fully oxidizing conditions, in the absence of respiratory substrates, and thus with an inactive respiratory chain. In mitochondria from statin-untreated astrocytes, CoQ10 supplementation increased both mtCoQ10H_2_ (∼8-fold) and mtCoQ10ox (∼25-fold, Fig. 7b) but not mtCoQ9H_2_ and mtCoQ9ox (Fig. 7a). As noted above, treatment with simvastatin alone lowered mtCoQ9H_2_ and mtCoQ9ox content by ∼40% and ∼25%, respectively, and mtCoQ10H_2_ and mtCoQ10ox content by ∼45% and ∼30%, respectively (Fig. 7a, b). In the mitochondria of statin-treated astrocytes, CoQ10 supplementation increased mtCoQ9ox content by ∼13%, but did not affect mtCoQ9H_2_ content, and increased mtCoQ10H_2_ and mtCoQ10ox content by ∼20% and ∼35%, respectively. The increased mtCoQ10H_2_ levels following CoQ10 supplementation in mitochondria from both statin-treated and untreated astrocytes indicates a substantial expansion of the antioxidant fraction of the mtCoQ pool.

CoQ10 supplementation did not alter maximal respiration under phosphorylating conditions during oxidation of the combined CI and CII substrates in mitochondria from statin-untreated astrocytes, but restored this oxidation in mitochondria of statin-treated astrocytes (Fig. 7c). Thus, CoQ10 supplementation reverted the statin-induced reduction in mitochondrial respiration.

During the oxidation of the combined CI and CII substrates, CoQ10 supplementation significantly decreased mtROS production and mtCoQ reduction level in mitochondria isolated from both statin-untreated and -treated astrocytes (Fig. 7d, e). Coadministration of CoQ10 with simvastatin eliminated the statin-induced increase in mtROS production (Fig. 7d). Thus, the decrease in the mtCoQ reduction level after CoQ10 supplementation in mitochondria of statin-treated astrocytes was accompanied by a decrease in mtROS production during respiratory substrate oxidation (Fig. 7e).

Overall, these results demonstrate that in mitochondria of statin-treated astrocytes, astrocyte CoQ10 supplementation markedly increases mtCoQ levels and lowers the mtCoQ reduction level, reduces mtROS levels, and restores mitochondrial respiration, thereby counteracting statin-induced disturbances in energy metabolism.

## 4 Discussion

Previous observations by our group showed that treatment with atorvastatin at a concentration of 200 nM or simvastatin at a concentration of 200 nM caused a comparable decrease in cellular CoQ levels in astrocytes, without negatively affecting cell viability [31]. Consistent with these results, the present study showed that exposure to the same concentration of these two statins caused similar changes in the bioenergetic functioning of mitochondria isolated from treated astrocytes.

### 4.1 Statin-induced CoQ deficiency

Mitochondria isolated from rat astrocytes treated with 200 nM atorvastatin or simvastatin for 6 days showed significantly decreased mtCoQ content (∼30%–35%) compared with those from untreated control astrocytes (Fig. 1a). These findings are consistent with the known mechanism of action of statins as inhibitors of HMGCR, a key enzyme in the mevalonate pathway. Because this pathway supplies the isoprenoid side chain necessary for CoQ biosynthesis [1, 3], its inhibition is expected to disrupt endogenous CoQ production and, thereby, reduce mtCoQ content. Moreover, statin treatment decreased COQ10B levels (Fig. 1b), suggesting that statins may affect not only the biosynthesis of CoQ but also proteins potentially involved in maintaining CoQ homeostasis within mitochondria [49]. Although the precise function of the COQ10B protein in mammals is not yet fully understood, it is proposed to be involved in the binding and intracellular transport of CoQ. Consequently, reduced expression of COQ10B may contribute to alterations in the availability and function of CoQ within mitochondria. However, further studies are needed to elucidate mechanisms underlying these effects, including the regulation of proteins essential for CoQ biosynthesis, such as COQ2, COQ7, or COQ9.

In our previous study, we showed an ∼35% decrease in total cellular CoQ9 and CoQ10 content, encompassing CoQ from all cell membranes, in astrocytes treated with atorvastatin or simvastatin [31]. In this study, we demonstrated for the first time that statins also reduce mtCoQ in astrocytes (Fig. 1). Crucially, only a few studies have directly demonstrated a statin-induced decrease in mtCoQ content [13, 14], as most prior studies have focused on total cellular CoQ levels. To date, mtCoQ10H_2_ levels following statin treatment have been reported primarily in muscle and liver cells [50, 51].

In this study, mtCoQ deficiency in statin-treated astrocytes was associated with a significant reduction in the antioxidant mtCoQH_2_ content, which represented ∼9% of mtCoQ9tot and ∼12% of mtCoQ10tot in untreated control astrocytes (Fig. 1a). Despite this decrease, we observed no changes in mitochondrial components of antioxidant defense, including SOD2 (Fig. 1b), nor in the levels and activity of energy-dissipating systems UCP and mitoBK_Ca_ (Fig. 6). These findings suggest that chronic statin exposure does not induce a mitochondrial antioxidant response, despite the increased mtROS production associated with the increased mtCoQ reduction (mtCoQH_2_/mtCoQtot) resulting from mtCoQ deficiency (Figs. 1 and 3). Interestingly, in our previous study, we showed decreased total ROS levels in statin-treated astrocytes at the cellular level but increased levels of cytosolic antioxidant enzymes, including superoxide dismutase 1 (SOD1) and glutathione-disulfide reductase (GSR) [31]. These observations suggest that compensatory antioxidant responses occur primarily at the cellular rather than the mitochondrial level.

### 4.2 Statin-induced changes in astrocyte mitochondrial function

The ∼35% statin-induced reduction in mtCoQ (mtCoQ9 + mtCoQ10) content (Fig. 1a) indicates a markedly lower availability of mtCoQ in the mitochondrial membrane, which may affect respiratory chain function, ATP synthesis, and mtROS production, thereby affecting overall energy metabolism. Mitochondria isolated from astrocytes treated with atorvastatin or simvastatin showed reduced maximal respiratory activity under phosphorylating conditions, decreased ΔΨ, and lower ATP synthesis compared with those from untreated control astrocytes (Fig. 2 and Table 1). Importantly, these bioenergetic changes were observed during both malate/pyruvate and succinate oxidation, indicating that respiration driven by both CI and CII is sensitive to statin-induced mtCoQ deficiency. This finding is consistent with a key role for mtCoQ as an electron carrier linking CI and CII to the downstream cytochrome pathway and suggests that mtCoQH_2_ availability impairs mitochondrial respiration supported by both CI and CII.

### Respiratory chain and OXPHOS

Reduced electron supply to the respiratory chain in the mitochondria of statin-treated astrocytes was associated with lower ATP production, as reflected by decreased ADP phosphorylation rates (Table 1) resulting from decreased respiration under phosphorylating state 3 conditions (Fig. 2a). These results are consistent with our previous study demonstrating a significant decrease in intracellular ATP content in astrocytes exposed to atorvastatin or simvastatin [31]. These data suggest that, under statin treatment, astrocytes may prioritize limiting mitochondrial ROS production and oxidative damage over maintaining ATP levels. This metabolic shift is consistent with the established role of astrocytes in maintaining redox homeostasis in the CNS [19, 52]. Unlike neurons, astrocytes rely more on glycolytic metabolism and are therefore less dependent on OXPHOS for ATP synthesis. This metabolic flexibility likely allows astrocytes to tolerate reduced mitochondrial ATP production while maintaining their antioxidant capacity and supporting the redox balance of the brain microenvironment. Consequently, statin-induced inhibition of mitochondrial electron transport may represent an adaptive response that minimizes oxidative stress at the expense of bioenergetic capacity. Interestingly, in the mitochondria of statin-treated astrocytes, the significant reduction in mtCoQ content (Fig. 1a) was accompanied by decreased activity and protein levels of the mtCoQ reducing complexes CI and CII (Fig. 5), as well as the mtCoQH_2_-oxidizing pathway CIII + CIV (Figs. 4 and 5). These changes may represent an adaptive response that limits both electron influx and efflux to/from the decreased mtCoQ pool under statin exposure. Moreover, unchanged UCP and mitoBK_Ca_ activities suggest that mitochondrial energy-dissipating systems do not decrease OXPHOS efficiency, as the ADP/O ratio remains unchanged.

The decreased activities of CI, CIII, and CIV observed in intact mitochondria from statin-treated astrocytes (Figs. 4 and 5) were associated with changes in the organization of respiratory supercomplexes, including a decrease in the activity and abundance of the CI+CIII_2_+CIV and CIII_2_+CIV supercomplexes (Fig. 5b). These structural rearrangements may represent an adaptive mechanism that limits mtROS production at mtCoQ-dependent sites, specifically the mtCoQ-reducing site in CI (I_Q_) [8, 9] and the mtCoQH_2_-oxidizing site in CIII (IIIQ_o_) [10]. Thus, remodeling of OXPHOS supercomplexes under mtCoQ deficiency may help reduce electron leakage and oxidative stress despite impaired respiratory activity. The molecular organization of the OXPHOS system has previously been reported to change in endothelial mitochondria under mtCoQ deficiency induced by statins, bisphosphonates, or hypoxia [15, 32, 33], suggesting that supercomplex remodeling may be a common mitochondrial adaptation to decreased mtCoQ availability.

To our knowledge, this study is the first to demonstrate that statins decrease the mitochondrial *a*-heme (cytochromes *a* + *a_3_*) content (Fig. 4b), potentially impairing mitochondrial respiration. The decrease in cytochromes *a* + *a_3_* content was accompanied by decreased reduction of these cytochromes (Fig. 4c), as well as diminished CIV activity in both intact mitochondria (Fig. 4a) and respiratory supercomplexes CI+CIII_2_+CIV and CIII_2_+CIV (Fig. 5b). Interestingly, although statins are known to inhibit the synthesis of the isoprenoid side chain required for *a*-heme biosynthesis [13, 14], a key component of COX (CIV), this effect has not, to our knowledge, been demonstrated experimentally until now. The tested statins did not alter the levels of the other mitochondrial cytochromes, cytochromes *c* (*c* and *c*_1_) and *b* (Fig. 4b), confirming that only isoprenoid tail-containing cytochromes were sensitive to mevalonate pathway inhibitors.

Interestingly, studies in permeabilized murine C2C12 myoblasts and isolated bovine heart mitochondria showed that statin lactones, rather than their acid forms, inhibit the Q_o_ site of CIII [18]. Our results suggest that prolonged exposure of astrocytes to statins may also impair CIII activity at this site, as evidenced by the decreased reduction of cytochromes *c* + *c*_1_ observed in mitochondria isolated from simvastatin-treated astrocytes (Fig. 4c). Moreover, direct administration of atorvastatin or simvastatin to mitochondria isolated from control astrocytes and rat brains decreased the reduction of cytochromes *c* + *c*_1_ like the action of myxothiazol, a well-known inhibitor of the Q_o_ site of CIII (Supplementary Fig. S4). Altogether, these findings indicate that statins can inhibit electron transfer through the Q_o_ site of CIII in mitochondria from astrocytes and the brain independent of mevalonate pathway inhibition. Notably, the inhibitory effect was observed after direct exposure to the acid forms of atorvastatin and simvastatin, suggesting that, at least in these mitochondrial preparations, statins do not need to be converted to lactone forms to interfere with CIII function.

### mtCoQ reduction level and mtROS formation during respiratory substrate oxidation

Our results indicate that chronic treatment of astrocytes with atorvastatin or simvastatin alters both the functional activity and the structural organization of the mitochondrial electron transport chain. Furthermore, the statin-induced ∼35% decrease in mtCoQ content (Fig. 1a) was associated with greater mtCoQ reduction level (mtCoQH_2_/mtCoQtot) under both phosphorylating and nonphosphorylating conditions (Fig. 3b). This change in the redox state of the mtCoQ pool was associated with enhanced mtROS production, with the most pronounced effect observed during malate/pyruvate oxidation when the I_Q_ and IIIQ_o_ mtCoQ-dependent mtROS production sites are engaged (Fig. 3a). Given that statins inhibit CIII by acting at the Qo site (Supplementary Fig. S6b) [18], reverse electron transport (RET) may be one possible mechanism contributing to the increased mtROS production observed in the mitochondria isolated from statin-treated astrocytes. This possibility is supported by the observation that succinate oxidation induced a greater increase in H_2_O_2_ release in the absence of CI inhibitor than in its presence (Supplementary Fig. S2a). However, the increased H_2_O_2_ release observed in the presence of rotenone also suggest the involvement of RET-independent mechanisms, potentially including ROS production at CII. Moreover, the decreases in the mtCoQ pool (Fig. 1a) and the activity of the mtCoQH_2_-oxidizing pathway (CIII+CIV, Fig. 4a) may contribute to the overall increase in mtROS production by increasing the mtCoQ reduction level during substrate oxidation in mitochondria isolated from statin-treated astrocytes. Further studies are needed to clarify the relative contributions of these mechanisms and identify the specific sites responsible for increased mtROS production following statin exposure.

Our findings suggest that statin exposure disrupts mtCoQ redox balance in astrocytes, not only by decreasing the mtCoQH_2_ antioxidant pool (nonoxidizable by the respiratory chain), but also by increasing the mtCoQ reduction level during respiratory substrate oxidation, which is an important factor in determining mtROS production [7, 45]. Notably, the statin-induced increase in the mtCoQ reduction level during respiratory substrate oxidation, along with the concomitant increase in mtROS production (Fig. 3), occurred without changes in the abundance or activity of antioxidant proteins, including SOD2, UCPs, and mitoBK_Ca_ (Figs. 1 and 6). Notably, this study is the first to demonstrate an association among alterations in mtCoQ content, mtCoQ reduction level, and mtROS production in the mitochondria of statin-exposed astrocytes.

### 4.3 Effect of CoQ10 supplementation

Our results indicate that supplementation with 3.4 µM CoQ10 significantly increases mtCoQ content (Fig. 7a, b) in the mitochondria of simvastatin-treated astrocytes. Moreover, CoQ10 restored mitochondrial maximal phosphorylating respiration impaired by statin treatment while simultaneously reducing mtROS production (Fig. 7c, d). The decrease in the mtROS production after CoQ10 supplementation in the respiring mitochondria of statin-treated astrocytes was accompanied by a significant decrease in the mtCoQ reduction level (Fig. 7e), further supporting the association between these two parameters.

These results indicate that CoQ10 supplementation ameliorates mitochondrial dysfunction and reduces mitochondrial oxidative stress in statin-treated astrocytes. Our previous cellular study of statin-treated astrocytes showed that CoQ10 supplementation increased intracellular CoQ10 content and reversed the statin-induced reduction in ATP levels [31]. Interestingly, although CoQ10 improved cellular bioenergetics, it did not further reduce overall cellular and mitochondrial ROS levels, suggesting that its beneficial effects are primarily related to maintaining mitochondrial energy metabolism rather than additional antioxidant activity at the cellular level. However, in this study, CoQ10 supplementation also attenuated the statin-induced increase in mtROS production in isolated mitochondria, suggesting a direct effect on mitochondrial redox homeostasis. This effect is likely related to the restoration of the CoQ10 pool in mitochondria and improved electron transfer in the respiratory chain, thereby reducing electron leakage and mtROS production. Thus, the marked increase in the oxidized mtCoQ10 pool following supplementation likely enhances the capacity of the mtCoQ pool to undergo redox cycling, rather than simply leading to an accumulation of potentially pro-oxidant quinone forms. CoQ deficiency can impair electron transport, promoting the buildup of semiquinone intermediates that may react with molecular oxygen to generate superoxide radicals. Replenishing the mtCoQ pool through CoQ10 supplementation can mitigate these disturbances, thereby improving electron transport and limiting the mtROS production. Altogether, our findings suggest that CoQ10 supplementation not only supports ATP synthesis but also helps maintain mitochondrial function and redox balance under statin-induced mitochondrial stress. These dual bioenergetic and redox-protective effects may contribute to CoQ10’s beneficial effects in statin-exposed astrocytes.

### 4.4 Potential consequences for astrocyte function and CNS pathology

The mitochondrial changes observed in statin-treated astrocytes may have important implications for CNS function and pathology. Astrocytes play a crucial role in maintaining brain redox homeostasis by providing metabolic support to neurons, regulating neurotransmitter recycling, and controlling neuroinflammatory responses [19–23]. Although astrocytes are more glycolytic and less dependent on OXPHOS than neurons, mitochondrial integrity remains essential for regulating ROS production, calcium homeostasis, and metabolic signaling [23]. Our findings demonstrate that statin-induced mtCoQ deficiency disrupts the mtCoQ redox balance, increases mtROS production, and impairs mitochondrial respiratory chain organization and function. Such changes may potentially affect the functioning of astrocytes and the support they provide to neurons, especially under conditions of increased metabolic demand. Mitochondrial dysfunction and oxidative stress in astrocytes are linked to neurodegenerative diseases such as Alzheimer’s disease, Parkinson’s disease, and amyotrophic lateral sclerosis [53–56].

Importantly, however, the moderate changes in mitochondrial bioenergetics induced by statins at a concentration of 200 nM were not associated with a decrease in astrocyte viability, as we demonstrated in our previous study [31]. Furthermore, in this study, we demonstrated that neither atorvastatin nor simvastatin affected glutamate uptake (Supplementary Fig. S7a), total nuclear factor kappa-light-chain-enhancer of activated B cells (NF-κB) protein levels, or the ratio of phosphorylated NF-κB to total NF-κB (Supplementary Fig. S7b) in astrocytes. These observations indicate that the capacity for glutamate uptake is preserved and that there is no detectable activation of NF-κB-dependent inflammatory signaling pathways. These results suggest that mitochondrial alterations induced by statins may represent metabolic adaptations that allow astrocytes to maintain key cellular functions despite limited mtCoQ availability and reduced overall OXPHOS capacity. Nevertheless, potential long-term consequences cannot be excluded, especially under conditions of increased metabolic demand or in the context of interactions between astrocytes and neurons. The ability of CoQ10 supplementation to improve mitochondrial function and attenuate oxidative stress (Fig. 7d) suggests that maintaining mtCoQ homeostasis may help preserve astrocyte metabolic function during prolonged statin exposure, which is associated with impaired mevalonate pathway activity. Further research using in vivo models is necessary to determine the physiological and pathological significance of these mitochondrial alterations and to assess their potential consequences for the functioning of CNS.

## 5 Conclusions

Prolonged exposure of astrocytes to atorvastatin or simvastatin disrupts mtCoQ homeostasis, leading to decreased mtCoQ content, increased mtCoQ reduction level associated with increased mtROS production in respiring mitochondria, and impaired mitochondrial bioenergetics. Statin-induced mtCoQ deficiency impairs mitochondrial bioenergetics by limiting electron transport, particularly through mtCoQ-related complexes. These changes are associated with reduced respiratory chain activity, reorganization of respiratory supercomplexes, decreased ΔΨ, and decreased ATP synthesis. Crucially, despite increased mtROS production, mitochondrial antioxidant and energy-dissipating systems remain unchanged, indicating that mitochondria in astrocytes do not elicit a compensatory stress response under prolonged statin exposure. Moreover, the observed mitochondrial changes do not appear to impair astrocyte function. CoQ10 supplementation increased mtCoQ levels, improved mitochondrial respiration, and reduced excess mtROS production, demonstrating the importance of mtCoQ homeostasis for astrocyte mitochondrial function. Therefore, this study identifies mtCoQ redox imbalance as a primary mediator of statin-induced mitochondrial dysfunction in astrocytes and highlights that CoQ10 supplementation may be a potential means of counteracting these effects.

## 7 Author Contributions

KW, LG, and WJ, conceptualized and designed the study; KW, LG, AB, and WP performed experiments: KW – main experimenter, all methods, AB, FG, and WP – assistance, and LG –

SDS-PAGE, BN-PAGE, in-gel activity, glutamate uptake; KW, LG, and WJ analyzed data; KW wrote the first draft of manuscript; WJ read the manuscript critically.

## 8 Acknowledgements

This research was funded by National Science Centre, Poland, OPUS 2020/37/B/NZ1/01188.

## Abbreviations

CNS: central nervous system
CoQ: coenzyme Q
mtCoQ: mitochondrial CoQ
mitoBKCa: mitochondrial big-conductance Ca2+-regulated potassium channel
mtCoQ: mitochondrial CoQ
mtROS: mitochondrial reactive species
OXPHOS: oxidative phosphorylation
RET: reverse electron transport
UCP: uncoupling protein
CI-CIV: respiratory chain complexes I-IV
ΔΨ: membrane potential

