## Supplementary figures for "Statin-induced Mitochondrial Coenzyme Q Deficiency Alters Mitochondrial Redox Homeostasis and Bioenergetic Function in Astrocytes"

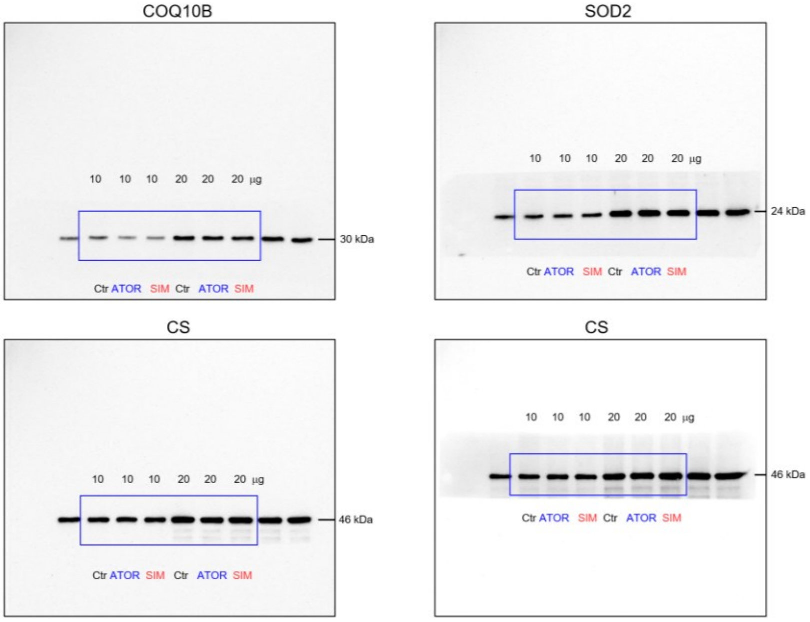


**Fig. S1.** Original images used to prepare Fig. 1. Abbreviations: Ctr, control; ATOR, atorvastatin; SIM, simvastatin; COQ10B, coenzyme Q-binding protein CoQ10 homolog B; CS, citrate synthase; SOD2, superoxide dismutase 2.

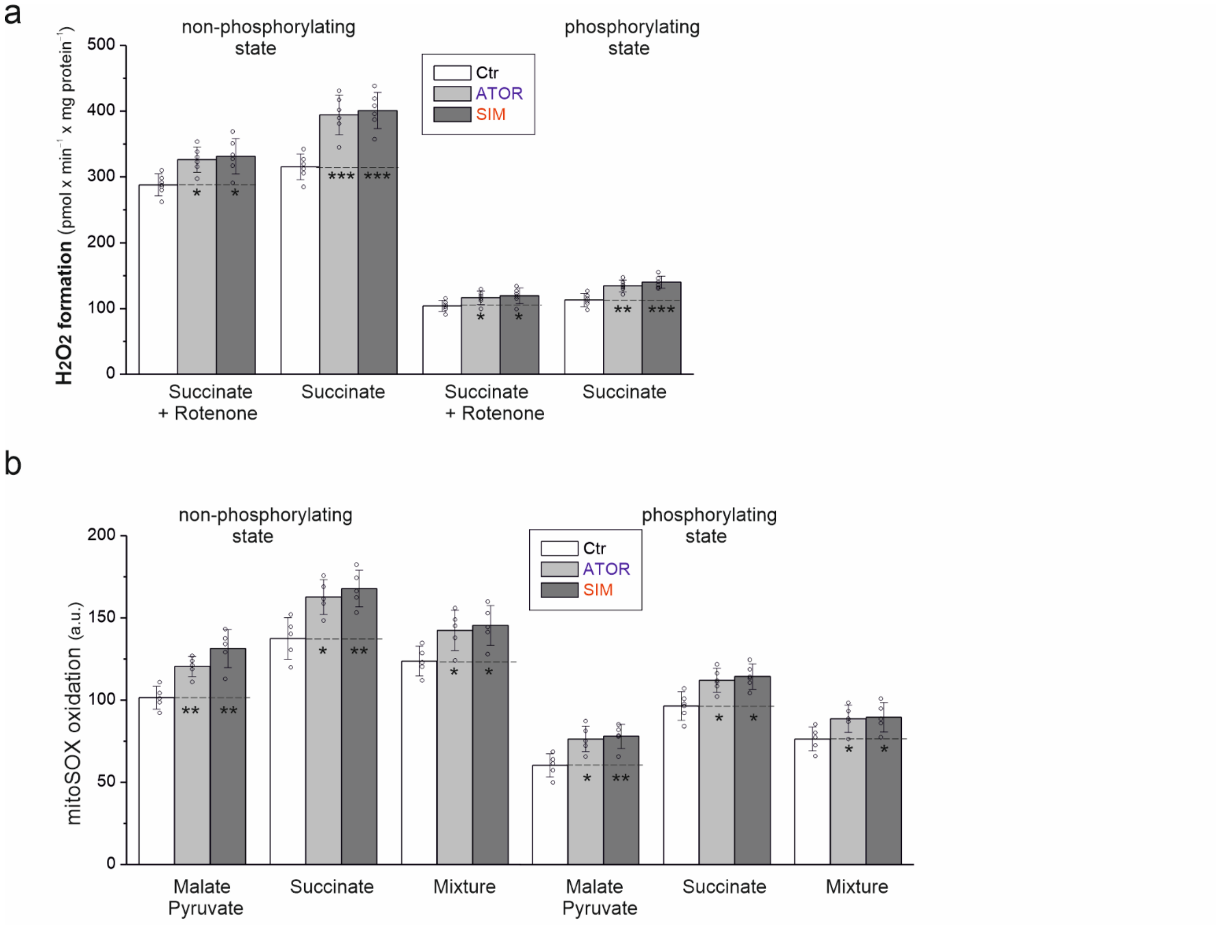


**Fig. S2**. **a**, $H_2O_2$ release in mitochondria isolated from untreated control astrocytes (Ctr) and atorvastatin (ATOR)- or simvastatin (SIM)-treated astrocytes during the oxidation of succinate in the absence or presence of rotenone under non-phosphorylating and phosphorylating conditions. **b**) Mitochondrial superoxide anion formation during the oxidation of various respiratory substrates under non-phosphorylating and phosphorylating conditions. **a**, **b**, Data are presented as the mean ± SD ($n$ = 5-6). Significance: *, $P$ <0.05; **, $P$ <0.01 compared with Ctrl (horizontal lines).

**b**) Mitochondrial superoxide anion formation was determined using a 2.5 µM MitoSox Red probe. Mitochondria (0.5 mg protein/ml) were incubated with fluorescent probe in a standard incubation medium (10 mM Tris-HCl [pH 7.2], 75 mM sucrose, 225 mM mannitol, 5 mM potassium chloride [KCl], 4 mM magnesium chloride [$MgCl_2$], 5 mM monopotassium phosphate [$KH_2PO_4$], and 0.15 mM EGTA). Mitochondrial protein samples were incubated with various respiratory substrates: 3 mM malate + 3 mM pyruvate, 3 mM succinate, or 3 mM malate + 3 mM pyruvate + 3 mM succinate for 10 min at 37°C. Assays were performed in the presence or absence of 1.5 mM ADP to assess mitochondrial ROS (mtROS) production under phosphorylating and non-phosphorylating conditions, respectively. The mitochondria were then washed twice with phosphate-buffered saline (PBS), centrifuged (12,200 × $g$ for 10 min at 4°C) and resuspended in PBS to a concentration of 0.5 mg protein/ml. Measurements were performed in 96-well plates with MitoSox (Excitation/Emission at 510/595 nm) using a Tecan Spark multiplate reader (Tecan Group Ltd). mtROS formation was expressed relative to control mitochondria isolated from astrocytes that had not been exposed to statins.

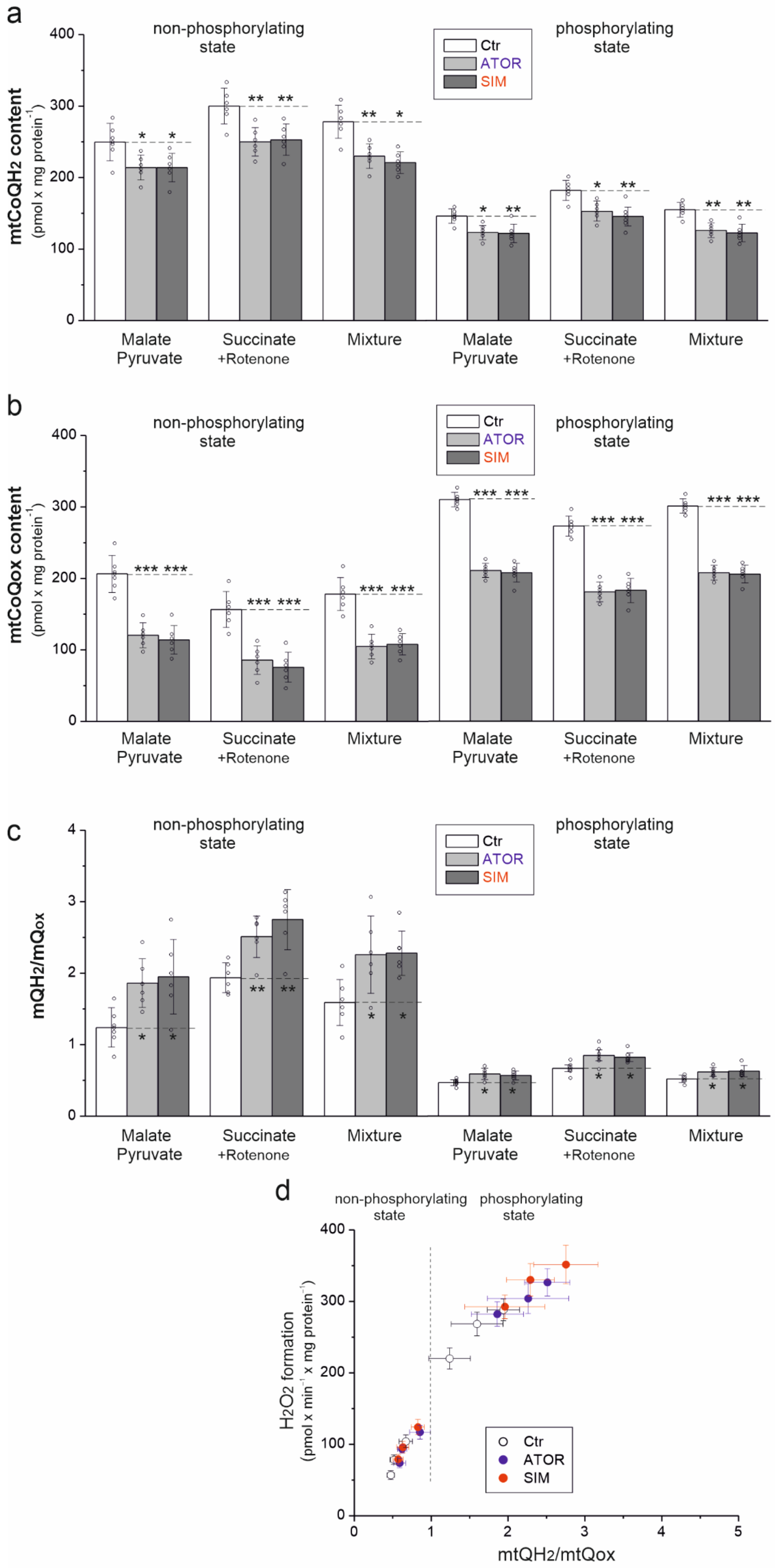


**Fig. S3**. (**a**) mtCoQH$_2$ content, (**b**) mtCoQox content mtCoQ, and (**c**) mtCoQ redox state (mtCoQH$_2$/mtCoQox), (**d**) relationships between $H_2O_2$ production and mtCoQH$_2$/mtCoQox of mitochondria isolated from untreated control astrocytes (Ctr) and atorvastatin (ATOR)- or simvastatin (SIM)-treated astrocytes during the oxidation of various respiratory substrates under non-phosphorylating and phosphorylating conditions. **a-d**, Data are presented as the mean ± SD ($n$ = 6). Significance (**a-c**): *, $P$ <0.05; **, $P$ <0.01; ***, $P$ <0.001 compared with Ctrl (horizontal lines).

a

b

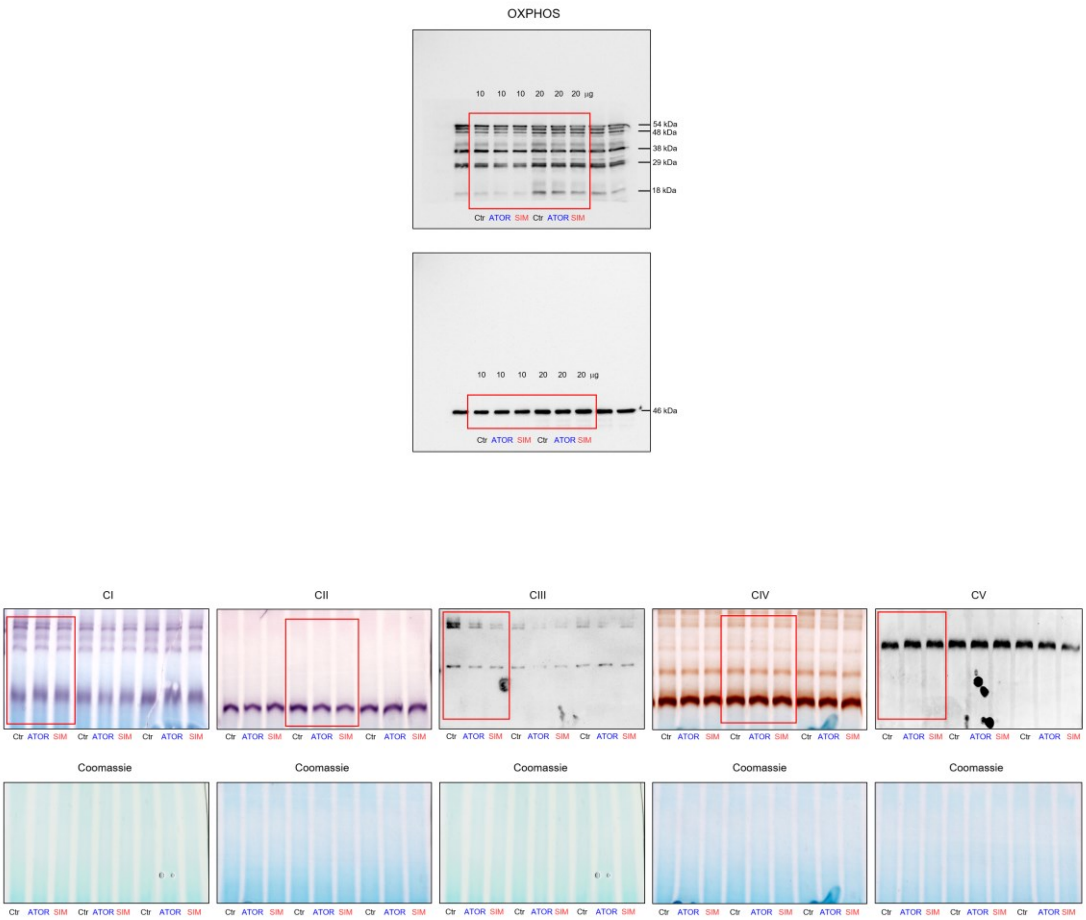


**Fig S4.** Source images utilized to generate Fig. 5. **a**, Full-length blots displayed in Fig. 5a. **b**, Gels used to prepare Fig. 5b, representing three biological replicates of respiratory complex activities or levels (top). The bottom panel displays the total protein loading control stained with Coomassie Brilliant Blue G-250 prior to activity staining. Red boxes indicate the specific lanes featured in the main figure. Abbreviations: Ctr, control; ATOR, atorvastatin; SIM, simvastatin.

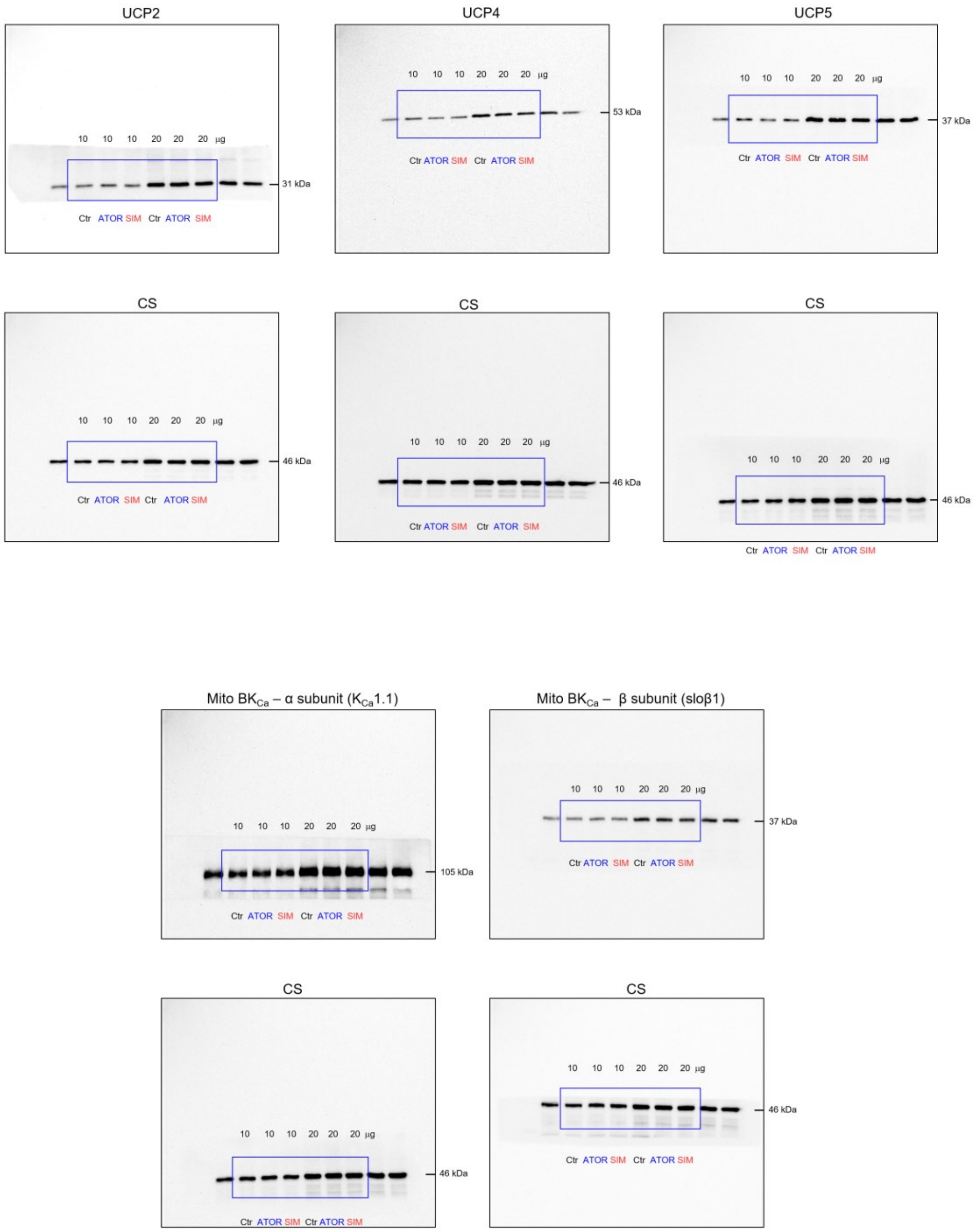


**Fig. S5**. Original images used for the preparation of Fig. 6c. Abbreviations: Ctr, control; ATOR, atorvastatin; SIM, simvastatin; UCP2, uncoupling protein 2; UCP4, uncoupling protein 4; UCP5, uncoupling protein 5; CS, citrate synthase; Mito $BK_{Ca}$, mitochondrial large-conductance $Ca^{2+}$-regulated potassium channel;

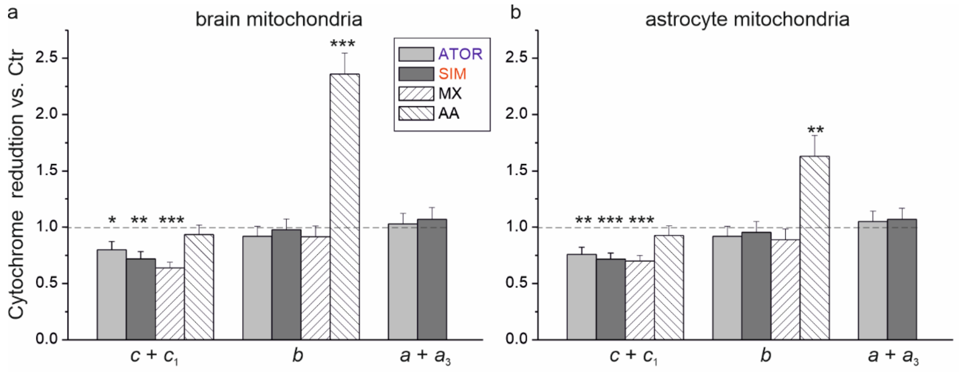


**Fig. S6.** Cytochrome reduction level in the mitochondria isolated from rat brains and untreated astrocytes. Mitochondria from rat brains were isolated as previously described [34]. Mitochondria were treated in vitro with 200 nM atorvastatin (ATOR), 200 nM simvastatin (SIM), 100 µM myxothiazol, or 30 µM antimycin A. Mean ± SD ($n$ = 5); $P$ <0.05 (*); $P$ <0.01 (**); P < 0.001 (***) relative to control mitochondria (horizontal lines)

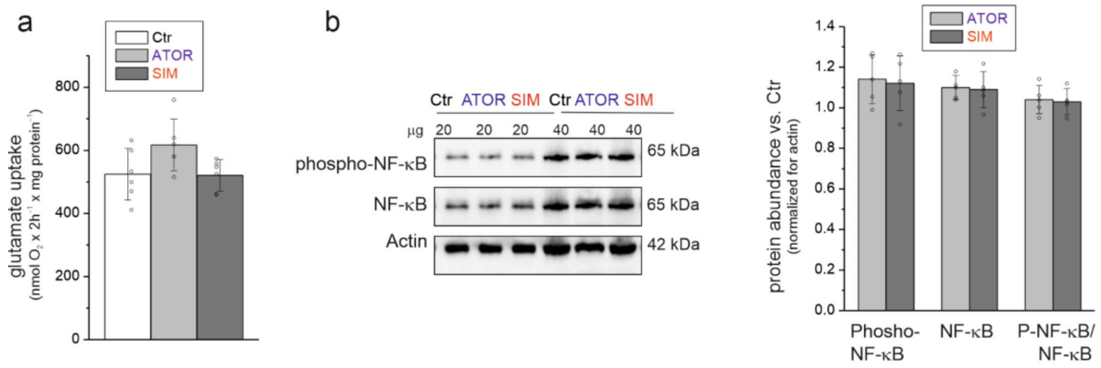


**Fig. S7**. Glutamate uptake efficiency (**a**) and the level of the inflammatory marker NF-κB (**b**) in untreated control astrocytes (Ctr) and in astrocytes treated with atorvastatin (ATOR) or simvastatin (SIM). **b**, Representative immunoblot images and quantification of protein levels. Loading control: actin. **a**, **b**, Data are presented as the mean ± SD ($n$ = 5-6).

*Glutamate uptake assay*

To assess the efficiency of glutamate uptake, after six days of culturing astrocytes (both control cells and those treated with statins) in 24well plates, the complete DMEM medium was replaced with serum-free and supplement-free DMEM medium containing 200 µM glutamate. After a 2-hour incubation period, the concentration of remaining glutamate in the culture medium was determined in a transparent 96-well plate using a commercial glutamate assay kit (catalog no. MAK330, Sigma-Aldrich, St. Louis, MO, USA), strictly following the manufacturer's instructions. Absorbance at a wavelength of 565 nm was measured using a Tecan Spark microplate reader (Tecan Group Ltd., Männedorf, Switzerland). The decrease in glutamate concentration was determined based on a standard curve. Values were normalized to the amount of cellular protein per well. To quantify the cellular protein, the cells used in the analysis were washed twice with phosphate-buffered saline (PBS), lysed using 100 mM NaOH, and the total protein content was then measured using the Bradford assay.

*Immunodetection of NF-κB levels*

The phosphorylation level of the p65 subunit of nuclear factor kappa-light-chain-enhancer of activated B cells (NF-κB) at Ser536 was determined by western blotting using an anti-phospho-NF-κB p65 (Ser536) antibody (catalog no. 3033; Cell Signaling Technology, Danvers, MA, USA) and normalized to total NF-κB p65 levels (anti-NF-κB p65 antibody, catalog no. 6956; Cell Signaling Technology, Danvers, MA, USA) and actin (42 kDa) (CP01, Calbiochem, San Diego, CA, USA).
